# *Tbx1* stabilizes differentiation of the cardiopharyngeal mesoderm and drives morphogenesis in the pharyngeal apparatus

**DOI:** 10.1101/2024.07.16.603669

**Authors:** Olga Lanzetta, Marchesa Bilio, Johannes Liebig, Katharina Jechow, Foo Wei Ten, Rosa Ferrentino, Ilaria Aurigemma, Elizabeth Illingworth, Christian Conrad, Soeren Lukassen, Claudia Angelini, Antonio Baldini

## Abstract

**Background:** TBX1 is required for the development of the pharyngeal apparatus. In the mouse, fish, and ascidian, *Tbx1* is a marker of cardiopharyngeal mesoderm (CPM), a cell population that provides progenitors to the heart and branchiomeric muscles. However, in mammals: a) the molecular cascade that drives the diversification of this multipotent cell population, and b) the role of *Tbx1* therein, are not well defined.

**Material and methods:** We used *in vitro* differentiation of WT and *Tbx1*^−/−^ mouse embryonic stem cells into precardiac mesoderm, and performed single cell RNA-seq and ATAC-seq at two differentiation stages. We then used WT and *Tbx1*^−/−^ mouse embryos for *in vivo* validation of the key findings.

**Results and conclusions:** We found that the response to loss of TBX1 is cell sub-population-specific, both in terms of gene expression and chromatin remodeling. We show that *Tbx1* regulates chromatin accessibility and gene expression of an ancient transcriptional module that orchestrates the development of the trunk, pharynx and heart across evolution. This module is co-regulated and includes genes encoding the conserved transcription factor families of Tea Shirt (Tshz), Sine Oculis (Six), Eye absent (Eya), and Ebf/Collier. Analysis of putative regulatory regions of these genes, which were selected using a machine-learning computational procedure, predicted a feed-forward regulatory relationship between TBX1 and SIX factors that drives or stabilizes the module. Most surprisingly, we found a drift in the differentiation trajectory of the *Tbx1* mutant CPM that led to a relative expansion of cells with epithelial-like transcriptional features in the cell culture model and in mouse embryos. We conclude that TBX1 is a critical factor for maintaining the transcriptional profile of the CPM.

## INTRODUCTION

Cardiopharyngeal mesoderm was defined as a cell population that is specified early in gastrulation that makes an important contribution to cardiac and craniofacial development^1,2^. Given that abnormal development of these structures leads to some of the most common birth defects such as congenital heart disease, understanding the mechanisms governing the developmental trajectory of this lineage has biological and clinical importance. Specifically, the molecular events that direct the differentiation of this population of multipotent mesodermal cells into more specialized cell types, chiefly skeletal and cardiac muscle, are not completely clear. Detailed studies in the ascidian *Ciona intestinalis* have defined a genetic map that guides their differentiation into pharyngeal and cardiac muscles^3,4^. In particular, these studies identified a critical node that takes a binary decision as to whether a cell will become pharyngeal or cardiac muscle in early development. A key factor in this decision is the transcriptional activity of TBX1/10, the homolog of the mammalian TBX1 protein, encoded by the gene *Tbx1*, the reduced dosage of which is causes DiGeorge syndrome^5–7^. *Tbx1/10* represses the cardiac fate of progenitors and triggers the pharyngeal fate^3^. Whether or not this activity is conserved in mammals is unclear.

Here, we used a cell differentiation model to address this key question about cardiopharyngeal mesoderm development. We found that *Tbx1* is “connected” to conserved transcription factors encoding genes that are known to play developmental roles in C. intestinalis, flies, and mammalian studies, and it supports both cardiac and pharyngeal expression programs not clear. We also found that loss of TBX1 causes a drift in differentiation that confers new properties to mutant cells.

We propose that TBX1 is required for the maintenance of an ancient transcriptional module that suppresses an epithelial-like drift of a subset of cardiopharyngeal progenitors.

## RESULTS

### A cellular model for cardiopharyngeal mesoderm cell diversification

We have used a serum-free mouse embryonic stem cell (mESC) differentiation protocol based on spontaneous aggregation, induction of mesoderm through transient exposure to Activin and BMP4, and then spontaneous differentiation into embryoid bodies (EB)^8^ (scheme in Fig. 1a). With this protocol, *Tbx1* gene expression is robustly activated between differentiation day 4 (d4) and d6 and continues through d8 (Fig. 1b), the last day that was tested. We next proceeded with single cell (sc)RNA-seq combined with scATAC-seq using the protocol suggested by the kit manufacturer (10xGenomics). We sequenced 8 samples at d6 and d8 for WT (parental cell line) and *Tbx1*^−/−^ mouse ES cells^9^ in two biological replicates (independent differentiation experiments). In total, we obtained high quality sequence of approx. 30000 cells. Cluster analysis, using integration of gene expression and ATAC data, identified 14 integrated clusters (ic) shown in Fig. 1c. The top 3 markers are shown in Fig. 1d, while a more extensive list of markers is reported in Suppl. Tab. 1.

**FIGURE 1.**
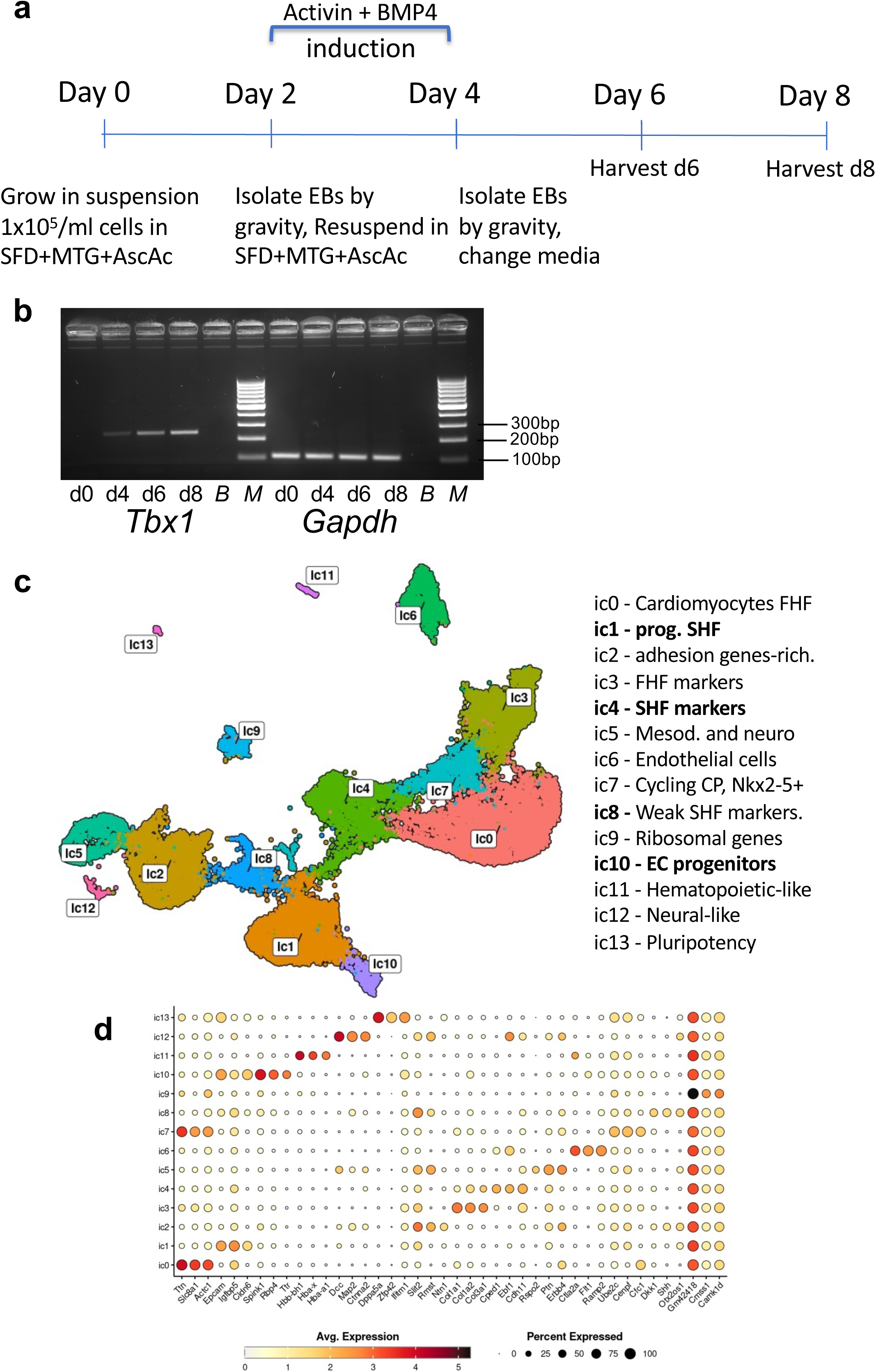
Description of a model for CPM differentiation from mouse embryonic stem cells. a) Cell differentiation scheme. SFD: serum-free differentiation made of 75% Iscove’s modified Dulbecco’s media (Cellgro Cat# 15-016-CV) and 25% HAM F12 media (Cellgro #10-080-CV). MTG: AscAc: ascorbic acid; EB: embryoid bodies. b) Reverse transcriptase-PCR assay of *Tbx1* gene expression and *Gapdh* (control) at d0-2-4-6-8 differentiation stages. c) Integrated (scRNAseq, scATACseq, d6, d8, WT, KO) datasets displayed on a uniform manifold approximation and projection (UMAP), separated into 14 clusters identified on the basis of marker gene expression. CM: cardiomyocytes; FHF: first heart field; SHF: second heart field; ECM: extra cellular matrix; CP: cardiac progenitors; EC: Endothelial cells. d) Dot-plot showing the expression of 3 top genes for each cluster. The full set of marker genes is reported on Supplementary Tab. 1.

Population-specific marker gene expression identified distinct clusters for the first heart field (FHF) and second heart field (SHF) lineages (Supp. Fig. 1a-b). Specifically, *Tbx1*, a marker of SHF and cardiopharyngeal mesoderm, was expressed in 5 clusters, including part of the endothelial cell cluster, ic6 (Supp. Fig. 1b, arrow and Suppl. Fig. 1c). Differentiation time affected cell distribution among clusters as visualized by the UMAP display, while the genotype had smaller impact on the overall cell population (Supp. Fig. 1d).

### Loss of Tbx1 has population-specific consequences within the CPM

We focused our attention on the cardiopharyngeal mesoderm clusters, as identified by SHF markers and *Tbx1* gene expression, i.e. clusters ic1, ic4, ic8, and ic10; cells assigned to these clusters were collected into a CPM subset (Fig. 2a-b) and reclustered using integrated RNA and ATAC data. This identified 5 clusters that are shown in Fig. 2b, in Fig. 2b’ is shown the same UMAP but color-coded with the same colors as in the integrated clusters shown in panel a (Fig. 2a). The top 3 marker genes of these clusters are indicated on Fig. 2c (additional markers are listed on Suppl. Tab. 1). The differentiation timepoint had a striking effect on clustering (Fig. 2d). Specifically, c0, c4, and part of c1 (indicated as c1/d6) were mostly represented at d6, while part of c1 (indicated as c1/d8), c2, and c3 were mostly represented at d8. The genotype also had a significant impact on cell distribution among clusters at d8: c2 was significantly expanded in the mutant while c3 was significantly reduced (Fig. 2e). Clusters included cardiac progenitors (markers *Kdr* and *Isl1*) (Suppl. Fig 2a), small populations of cardiomyocytes (*Ttn*) and smooth muscle cells (*Tagln*) (Suppl. Fig 2b-c), branchiomeric muscle progenitors (*Tlx1, Tcf21*) (Suppl. Fig 2d), and branchiomeric muscle cells (*Msc*, *Myf5*) (Suppl. Fig 2e). Of note, we did not detect posterior SHF cells in the CPM subset, as indicated by the lack of detectable expression of *Hoxb1* and *Tbx5* (Suppl. Fig. 2f-g)*. Tbx5* was expressed in the FHF domain, as expected (Suppl. Fig. 2h). We measured the Euclidean distance between the transcriptional profiles of the CPM clusters (after pseudo-bulk cumulation of data within each cluster) using multidimensional scaling (MDS); results identified high similarity between c0/d6 and c2/d8, and between c1/d6 and c3/d8 (Suppl. Fig. 3a), suggesting that between d6 and d8, c0 cells take on a c2 profile, while c1/d6 cells take on a c3 profile. This hypothetical “flow” is supported by a trajectory analysis using Monocle3^10^ (Suppl. Fig. 3b).

**FIGURE 2.**
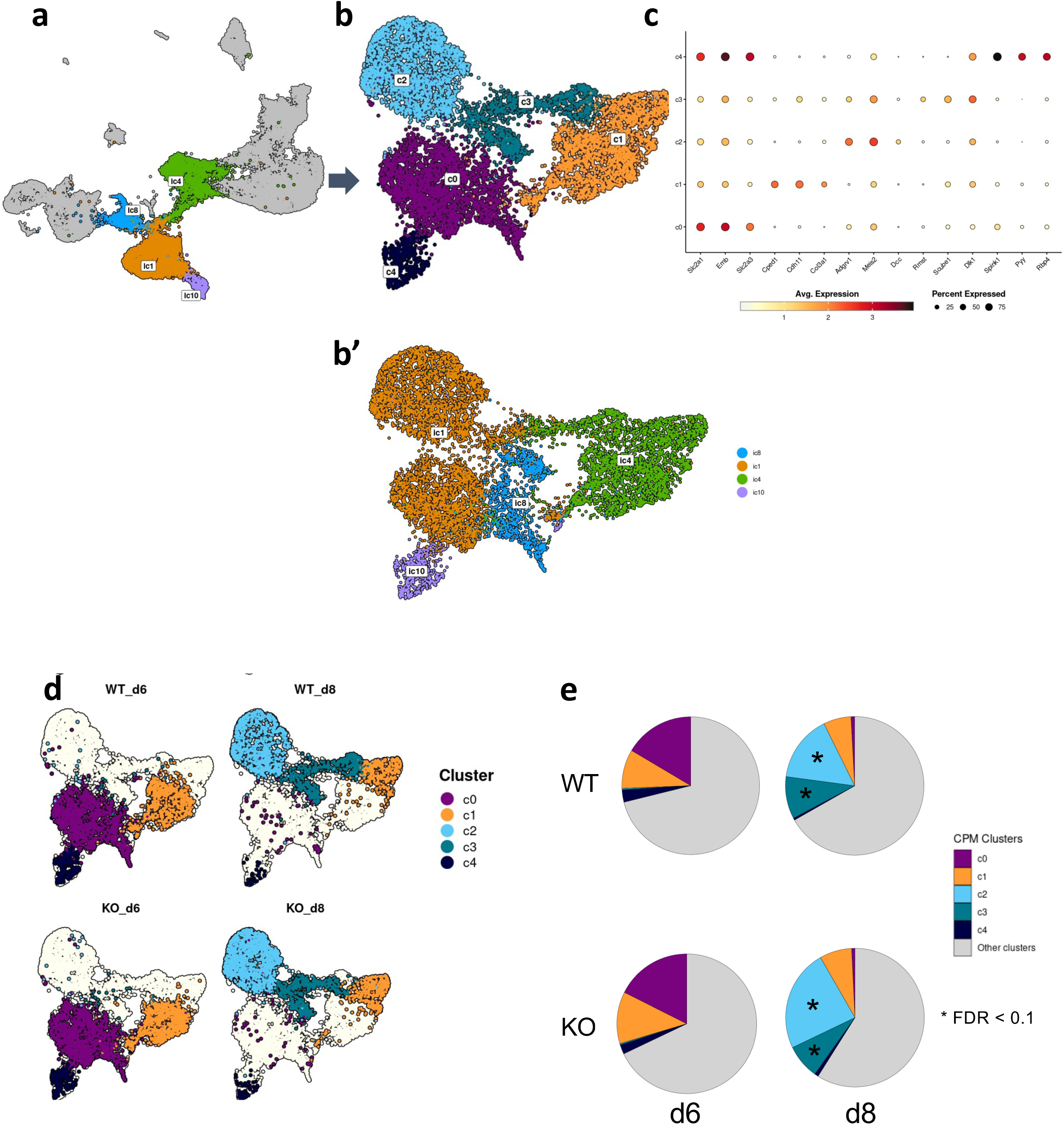
Re-clustering the CPM/SHF population. a-b) Clusters ic1, 4, 8, and 10 of the general population were re-clustered, using scRNA-seq and scATAC-seq data, into a CPM subset subdivided into 5 clusters named c0 to c4. b’) same UMAP as in (b) but color-coded to identify the re-location of cells from ic clusters 1, 4, 8, and 10. c) Top 3 markers of each cluster of the CPM subset. The full set of marker genes is reported in Supplementary Tab. 1. d) Clusters of the CPM subset are mostly d6- or d8-specific. e) Cell distribution is affected by genotype, c2 is over-represented in the KO dataset, while c1 is under-represented. Gray areas indicate the other clusters of the full dataset.

Loss of *Tbx1* affected chromatin accessibility in correspondence with previously published ChIP-seq data; 2388 TBX1 ChIP-seq peaks from early differentiating cells^11^ (Fig. 3a); even more evident was the effect on the 255 ChIP-seq regions identified in mouse embryos^12^ (Fig. 3b). The union of all of the differentially accessible regions (DARs) calculated in the CPM dataset at d6 and d8 identified 185 unique regions (Suppl. Tab. 2, examples of coverage of selected DARs in Suppl. Fig. 4); a search for enriched transcription factor motifs in these regions identified a T-box motif as the most enriched motif, and homeobox-binding motifs (Fig. 3c, full results are shown in Suppl. Tab. 3). A gene ontology (GO) search of genes neighboring DARs is shown in Suppl. Tab. 4. Overall, a large portion of chromatin changes identified in the mutant CPM dataset demonstrated reduced accessibility and may be directly attributed to the loss of TBX1. These results are also consistent with the proposed role of TBX1 in chromatin remodeling^13^. We also calculated differentially expressed genes for each cluster of the CPM subset and found transcriptional changes mostly mapped to c3, followed by c1 and c2 (genes listed in Suppl. Tab. 5).

**FIGURE 3.**
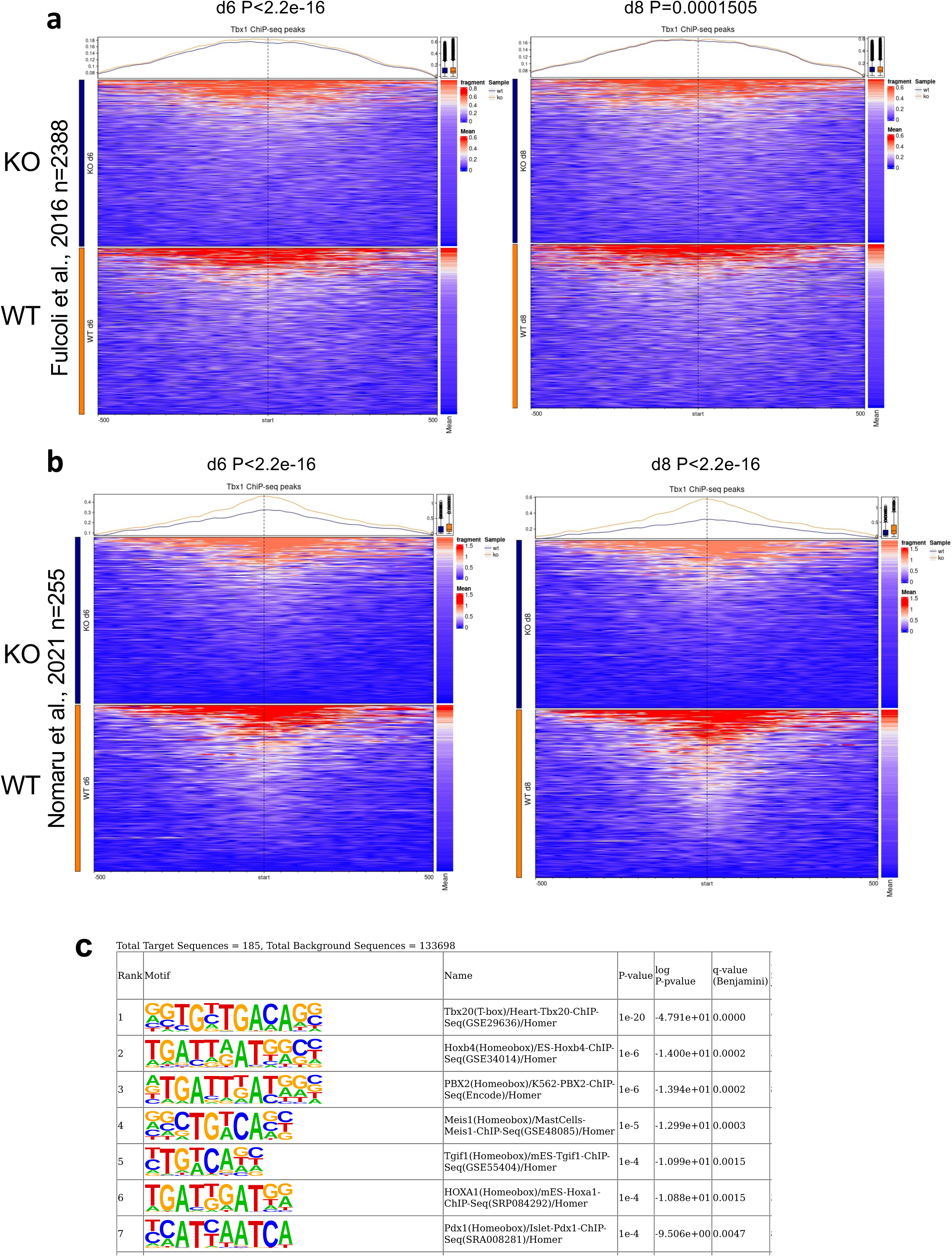
Reduced chromatin accessibility of Tbx1 binding sites in mutant cells. a) Heat maps showing chromatin accessibility at d6 (left panel) and d8 (right) of KO (top) and WT (bottom) cells of the CPM dataset of the TBX1 ChIP-seq peaks (n=2388) reported in^11^. At both time points, the coverage is significantly lower in KO cells. b) Heat maps showing chromatin accessibility at d6 (left panel) and d8 (right) of KO (top) and WT (bottom) cells of the CPM dataset of the TBX1 ChIP-seq peaks (n=255) reported in^12^. At both time points, the coverage is significantly lower in KO cells. Chromatin accessibility at Tbx1-ChIP regions (255 peaks reported in ^12^. Differences are statistically significant. c) Enriched binding motifs identified in the differentially accessible regions (n=185) found in the CPM subset. The top scoring motif is a T-box binding motif.

### Tbx1 is connected with and regulates transcriptional modules related to CPM cell diversification

To identify the gene sets most likely to drive diversification within the CPM, we used the weighted correlation network analysis (WGCNA) algorithm adapted to single cell data (high dimensional WGCNA)^14^. Results identified co-regulated gene-sets, hereafter referred to as transcriptional modules (Suppl. Tab. 6). We then used a differential module eigengene analysis to evaluate differences between WT and *Tbx1*^−/−^ cells for each module and identified modules that were responsive to the loss of *Tbx1* (Suppl. Tab. 6). We focused on c1 and c3 where there was the most prevalent *Tbx1* gene expression and transcriptional changes. In c1, the most affected module was c1mod4 (P= 1.82e-31), this module includes *Tbx1* and is enriched in genes encoding transcription and signaling factors involved in tissue morphogenesis, including genes involved in branchiomeric muscle development and cardiac development (Suppl. Tab. 6 and Fig. 4a, gene ontology search results are shown in Suppl. Tab. 4). The second most affected module was c1mod8 (P= 4.92e-11) (Fig. 4b), which is enriched in genes involved in cardiac development (Suppl. Tab. 4). Heat maps of expression of hub genes for the two modules are shown in Suppl. Fig. 5a-b. We next identified putative regulatory elements of c1mod4 and c1mod8 hub genes, defined as ATAC peaks that mapped within the genes or within 15Kb upstream of the transcription start site (TSS) or downstream of the transcription end site (TES), and with a positive score in the logistic regression test (score >0.5). These regions are listed in Suppl. Tab. 7. Chromatin accessibility data derived from the simultaneous scATAC-seq assay showed that loss of *Tbx1* was associated with significantly reduced accessibility of promoters and putative regulatory sequences (Fig. 4c,e for c1mod4 and Fig. 4d,f for c1mod8).

**FIGURE 4.**
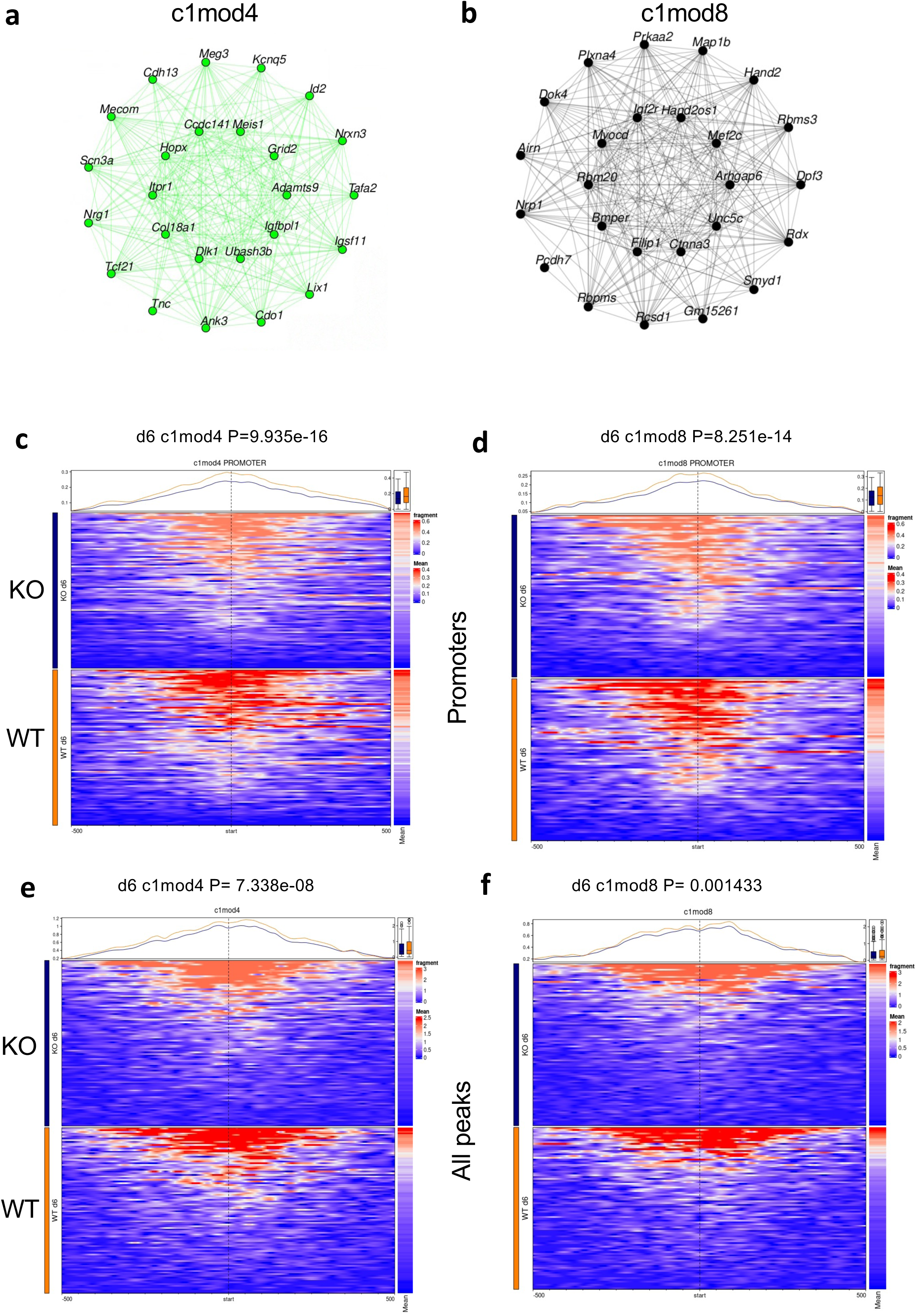
Transcriptional modules in the CPM subset: early time point. a-b) Connections among hub genes of modules c1mod4 and c1mod8. For each dendrogram, the inner circle is represented by genes that are more tightly connected, on the basis of co-expression evaluation. c-f) The chromatin state of hub genes of the two modules responds significantly to loss of the *Tbx1* gene at promoters (c-d), and over the entire gene body (e-f).

The expression of c1mod4 and c1mod8 genes generally straddles the c1/d6 and c3/d8 clusters, and some of those genes’ expression resembles the expression of *Tbx1,* further supporting the relatedness of the c1 and c3 transcriptomes, as suggested by multi-dimensional scaling and trajectory tests shown in Suppl. Fig. 3. For example, the expression of c1mod4 top scoring genes *Grid2* and *Adamts9* appeared partially overlapping with *Tbx1* in our WT CPM dataset at d6 and d8 (Fig. 5a-b). *In situ* hybridization of E9.5 mouse embryos confirmed the overlap of *Grid2* and *Tbx1* expression (Fig. 5c) and showed a substantial reduction of expression in *Tbx1*^−/−^ embryos (Fig. 5d-d’, arrows indicate the SHF region of the dorsal pericardial wall). *Adamts9*, which showed broader overlap with *Tbx1* at d6 and d8 (Fig. 5b), was also expressed in the first pharyngeal arch (PA-1) of embryos (arrows in Fig. 5e-e’) whereas in the mutant, the expression was virtually abolished in this domain, although it was still detectable in other domains (Fig. 5e-e’). Thus, on the whole, the two *Tbx1*-dependent modules of the c1/d6 cluster appear to be unrelated because one is enriched for cardiac development genes (c1mod8), while the other (c1mod4) is more heterogeneous and includes morphogenesis genes broadly related to the development of the pharyngeal apparatus.

**FIGURE 5.**
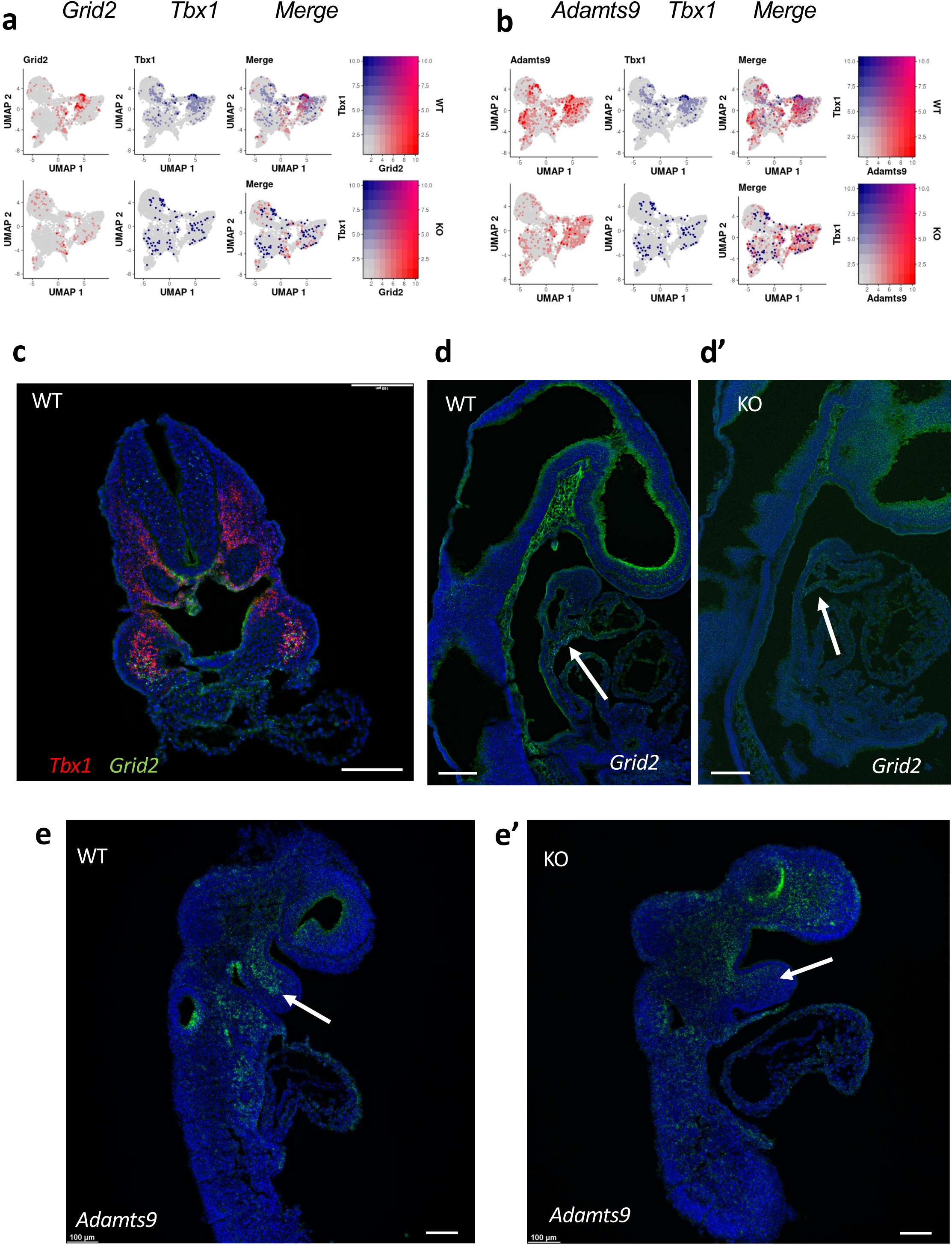
Top-scoring hub genes of c1mod4 Grid2 and Adamts9 are co-expressed with Tbx1 and are downregulated in KO cells in cell cultures and in vivo. a-b) The UMAPs indicate substantial overlap between *Grid2* (a) or *Adamts9* (b) and *Tbx1* expression. Note that the mutant allele of Tbx1 is lightly transcribed because the gene has been mutated by insertion rather than deleted, as described^9^. c) A transverse section of a WT E9.5 embryo stained by RNAscope using probes for *Tbx1* (red) and *Grid2* (green). Note the substantial, albeit not complete overlap, especially at the core the second pharyngeal arches. d-d’) Sagittal sections of WT (d) and KO (d’) moues E9.5 embryos stained by RNAscope using a *Grid2* probe, note the reduction of signal in the SHF (arrows) of the mutant embryo. e-e’) Sagittal sections of WT (e) and KO (e’) moues E9.5 embryos stained by RNAscope using an *Adamts9* probe. The signal in the core mesoderm of the first pharyngeal arch (arrows) is undetectable in the mutant embryo. Scale bars are 100 µm.

We next focused on module c3mod4, a d8 module that is down regulated in the mutant cell line and includes *Tbx1*. The module is enriched in genes involved in the development of the pharyngeal apparatus and, particularly, in the development of the branchiomeric muscles (Suppl. Fig. 6a; gene expression heat map in Suppl. 5c; hub genes in Suppl. Tab. 6; GO search in Suppl. Tab. 4). Therefore, it is related to c1mod4 with which it shares 20% of the hub genes. The module includes genes *Tbx1*-*Six*/*Eya*-*Ebf*, *Tcf21* that encodes a group of transcription factors that drives muscle differentiation by activating myogenic regulatory factors (MRF)-encoding genes (e.g. *Msc*, *Myf5*). These genes are shut down in mutant cells of the CPM subset (Suppl. Fig. 7) and their expression is altered in *Tbx1* mutant mouse embryos^15^. The pathway TBX1-EBF->MRF has already been described in the ascidian C. intestinalis as a driver of pharyngeal muscle development and differentiation^3^ while, in the mouse, interactions between *Tbx1*and *Six*/*Eya,* and between *Tbx1* and *Tcf21* have been reported^16,17^. The highest scoring gene in module c3mod4 was *Tshz2*, which encodes a Zinc-finger and homeobox transcription factor that functions as a homeotic protein that specifies trunk development in *Drosophila*^18,19^; morpholino-based suppression of the *Tshz2* Zebrafish homolog caused severe defects of craniofacial morphogenesis^20^ but its role in pharyngeal development in mammals has not been reported. We found that in E9.5 mouse embryos, the *Tshz2* expression pattern was similar to that of mesodermal *Tbx1* expression, mainly in the core mesoderm of the pharyngeal arches and in the dorsal pericardial wall (DPW), sites of derivatives of the CPM (Suppl. Fig. 8a). In *Tbx1*^−/−^ embryos, *Tshz2* expression was reduced overall and rearranged due to the anatomical changes that occur in this mutant and lead to the anomalous distribution of CPM cells; signal down regulation was particularly evident in the core mesoderm of PA-1 (arrows in Suppl. Fig. 8b). Thus, *Tshz2* expression, while correlated to *Tbx1* gene expression, was reduced but not abolished by the loss of TBX1 function. Chromatin accessibility of putative regulatory elements associated with c3mod4 genes was reduced in mutant cells (Suppl. Fig. 6b).

### The SIX family of transcription factors in c3mod4 are candidate mediators of the response to Tbx1

To identify candidate regulators of transcriptional modules, we used the putative regulatory elements of genes of c1mod4, c1mod8, and c3mod4 (Suppl. Tab. 6) and performed HOMER search using all the peaks (n=134191) in the dataset (peakome) as the background. Results showed no significant enrichment of binding motifs in c1 modules, possibly due to the paucity of sequences or cell heterogeneity. However, we found significant motif enrichment in c3mod4 regions (n=711), specifically we found PITX1/E-BOX, and NFAT-AP1 motifs (Suppl. Fig. 6c). To understand whether the c3mod4 subset of regulatory elements that are made less accessible by *Tbx1* loss of function are enriched in specific binding motifs, we selected peaks that were less open in the mutant cells (logFC < −log2(1.2)) (n=131) and ran HOMER again. We found the SIX1/SIX2 motif significantly enriched (Suppl. Fig. 6d), suggesting that this family of transcription factors (evolved from the fly *sine oculis*) are intermediaries of this module responsiveness to *Tbx1* loss of function. We did not find enrichment of T-BOX motifs, suggesting that TBX1 may function as an upstream primer of some of these factors, e.g. *Six1-4*, *Eya1-4* (encoding co-factors of SIX transcription factors), and *Tcf21* genes, as in fact suggested by ATAC-seq and gene expression data that show regions of reduced accessibility in the *Tcf21* and *Six1* genes and reduced expression of *Eya1* and *Eya4* in mutant cells at d8 (Suppl. Fig. 9).

### Loss of Tbx1 is associated with a drift in CPM differentiation

We used RNA velocity-based tests (implemented in DYNAMO^21^) to leverage transcriptional dynamics and infer cell differentiation progression within the CPM subset. Velocity display showed similar dynamics in WT and *Tbx1*^−/−^ cells, identifying 3 main trajectories indicated by yellow arrows in Fig. 6a-b: c0->c2, c1d6->c3, and c3->c2. The latter relation was not predicted by MDS and Monocle3 (see Suppl. Fig. 3a-b). In mutant cells there was a relative increase of the cell numerosity of c2 and relative decrease of c3, compared to WT cells (Fig. 2e), suggesting an enrichment of c2-like transcriptome and an impoverishment of the c3-like transcriptome in mutant cells. Gene expression analyses showed that c2 was enriched in epithelial markers, cell adhesion, and basement membrane related genes (GO search results in Suppl Tab. 4). We found at least two genes, *Postn* and *Otx2*, for which expression was expanded, and their transcriptional dynamics accelerated in mutant clusters (Fig. 6c, compare with control gene *Fn1*, panel d). In E9.5 WT mouse embryos, these genes are mainly expressed in epithelial tissues. *Postn* gene expression was expanded in c2 of *Tbx1* KO cells (arrow in Fig. 6e) and in the mesodermal core of PA-1 of E9.5 *Tbx1*^+/−^ and *Tbx1*^−/−^ embryos, compared to WT embryos (arrows in Fig. 6f). At this stage, *Postn* is mainly expressed in the pharyngeal ectoderm of the WT embryo (Fig. 6f, arrowhead). The accessibility of the *Postn* gene promoter region was also increased in mutant cells in c3 (Fig. 6g). *Otx2* gene expression was expanded in mutant c1 and c2 cells (Suppl. Fig. 10a, arrows), it exhibited increased opening of the promoter region in mutant c3 cells (Suppl. Fig. 10b), and was up regulated in the core mesoderm of PA-1 and head mesenchyme of *Tbx1*^−/−^ embryos at 16st and 26st stages (by immunofluorescence, arrowheads in Suppl. Fig. 10c, and by in situ hybridization, arrows in Suppl. Fig. 11). Double immunofluorescence with anti-OTX2 and anti-periostin antibodies revealed similar distribution of the two signals in the core mesoderm of the E9.5 *Tbx1*^−/−^ PA-1 (Fig. 7a). The periostin signal morphology in the core mesoderm resembled a basal membrane (magnified detail in Fig. 7a), further suggesting the epithelial-like character of these cells.

**FIGURE 6.**
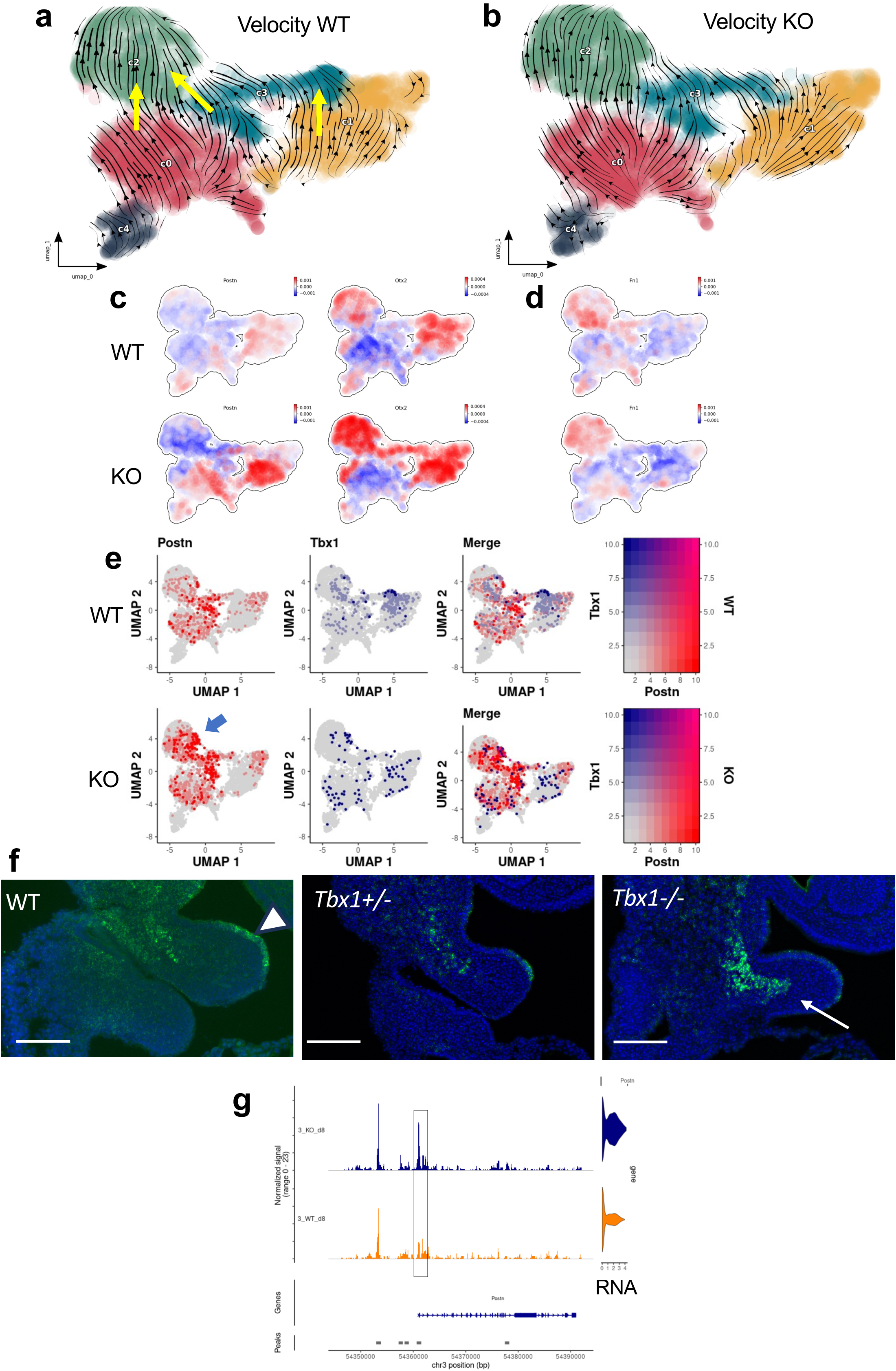
Loss of Tbx1 has an only minor effect on the differentiation trajectory but it up regulates inappropriate genes. a-b) Velocity maps of the WT (a) and mutant (b) CPM subset. There are three directions highlighted by yellow arrows: c0->c2, c3->c2, and c1->c2. The trajectories of WT and KO cells are very similar. c-d) Color map shows transcriptional acceleration of *Otx2* and *Postn* (c), compared to a control gene, *Fn1,* that is not significantly affected by loss of *Tbx1*(d). e) The *Postn* gene expression does not normally overlap with *Tbx1* gene expression (e, top panel), but it is expanded in the c2 domain (arrow, bottom panel) in the mutant. f) Sagittal sections of E9.5 mouse embryos of the indicated genotype, stained by RNAscope using a probe for *Postn*. Note the expansion of staining in the first pharyngeal arch of mutant embryos. Scale bars are 100 µm. g) ATAC coverage of the *Postn* gene in cluster c3 cells of the CPM subset. Note the increased opening of the promoter region in mutant cells.

**FIGURE 7.**
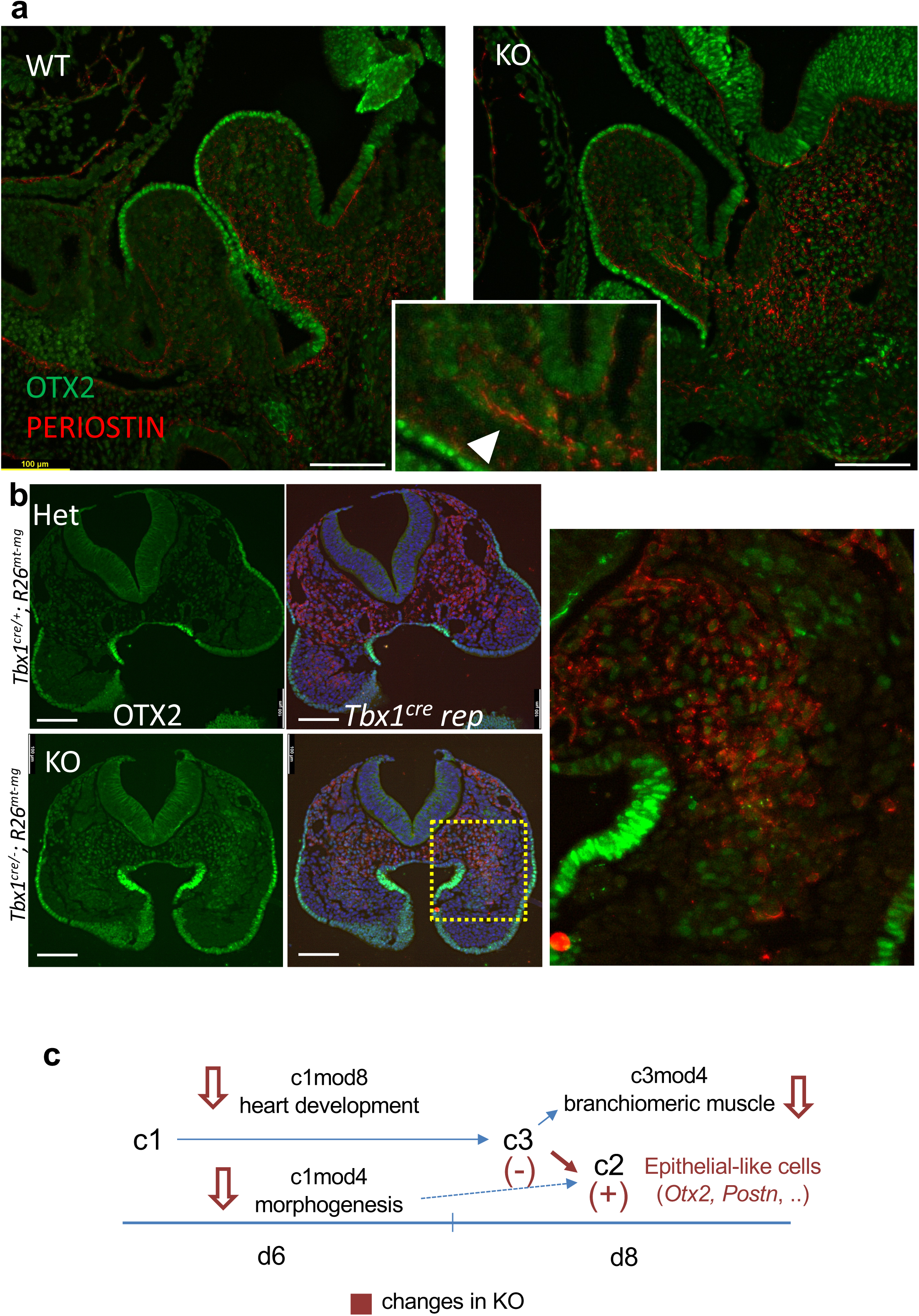
Up regulation of the Otx2 gene in the core mesoderm of PA-1 and the head mesenchyme in the mutant is in part a cell autonomous phenomenon. a) Double immunofluorescence OTX2-Periostin on mouse E9.5 sagittal sections, note the basal-membrane-like arrangement of periostin in the mutant PA-1 (arrowhead in the magnified inset). b) Transverse sections of E9.5 mouse embryos of the indicated genotype in which *Tbx1* expressing cells and their descendants are genetically labelled (in red). The green signal derives from immunofluorescence with an OTX2 antibody. Note the lack of overlap in the hetrozygous (control) embryo (top left panels) but extensive overlap in the homozygous mutant embryo (bottom panels, highlighted in the magnified inset). Images from additional genetic labelling experiments are reported on Supplementary Figs. 12 and 13. Scale bars are 100 µm. c) A cartoon summarizing our findings and our working hypothesis for future studies. Changes observed in mutant cells are indicated in red-brown. c1 includes at least two transcriptional networks relevant for CPM fate, pharyngeal morphogenesis (branchiomeric) and heart (SHF) development. c1 cells converge into c3 where they start their differentiation into branchiomeric muscle and, in part, cardiac progenitors. From c3, a group of cells assumes a c2-like transcriptional profile, characterized by ECM-related and cell adhesion gene expression. This phenomenon is more evident in mutant cells where a set of genes broadly related to epithelial cells is ectopically activated.

To understand whether the expression of *Otx2* in the core mesoderm of mutant embryos occurred in cells expressing *Tbx1* or in their descendants, i.e. if the up regulation of the *Otx2* gene is a cell autonomous event, we performed OTX2 immunofluorescence in *Tbx1^cre/+^*;R26^mT-mG^ (heterozygous control) and *Tbx1*^*cre*/−^;R26^mT-mG^ (null) E9.5 mouse embryos. In these genotypes, *Tbx1* expressing cells and their descendants are genetically labeled by GFP staining (revealed with anti-GFP antibodies). The *Tbx1^cre^* allele is functionally null but it expresses the Cre recombinase, therefore the strategy labels genetically *Tbx1*-expressing cells in the absence of functional TBX1. Results showed extensive overlap between the GFP and OTX2 signals in *Tbx1* null E9.5 mouse embryos (Fig. Fig. 7b, and additional examples in Supp. Figs. 12 and 13) indicating that at least part of the up regulation of OTX2 occurs in cells that normally express *Tbx1* and their descendants.

Collectively, these observations demonstrate that *Tbx1* is required to suppress a transcriptional drift towards an epithelial-like profile in the PA-1 and head mesenchyme.

## DISCUSSION

We used a cellular differentiation model to study the role of *Tbx1* in CPM diversification and differentiation using combined scRNA-seq and scATAC-seq on the same cells. We show that the differentiation protocol is suitable to obtain most of the various cell types described *in vivo* including, and most relevant for this study, cardiac and branchiomeric muscle progenitors, although we were unable to identify pSHF cells. The lack of a posterior program will be an interesting topic for further studies, as it suggests that the model lacks or is unable to respond to posteriorizing signals, such as retinoic acid^22^.

We tested two incubation time-points, d6 and d8, and already at d6, we could clearly distinguish FHF and SHF populations, the two major mesodermal lineages contributing to heart development. Smooth muscle cells seem to have diverged before our earliest time point. At d6, a multilineage program is evident, exemplified by the c1mod4 module, with heart progenitors and branchiomeric muscle progenitors. This module was strongly affected by loss of *Tbx1*. However, our data did not identify clearly a multi lineage primed (MLP) cell population as defined by Nomaru et al.^12^, although the c1/d6 cluster may be a close relative. The lack of a posteriorizing transcription program may have contributed to the difficulty in identifying MLP transcriptomes in our model because the main location of the MLP population is in the caudal region of the pharyngeal apparatus^12^. A branchiomeric muscle developmental module was clearly evident at d8, which was also strongly affected by loss of *Tbx1*, both in terms of chromatin landscape and gene expression.

We found that *Tbx1* is a component of an evolutionary conserved gene program, that we have named c3mod4, that includes genes involved in heart and branchiomeric muscle development, including *sine oculis*-like and its cofactors *eye absent*-like factors as well as EBF/Collier factors. Enhancer selection and binding motif searches suggest that *sine oculis* factors are among the transcription factors regulating the module downstream to *Tbx1*.

In the ascidian C. intestinalis*, Tbx1/10* is required to switch the fate of pluripotent cells towards a pharyngeal muscle fate rather than towards cardiac muscle, thus predicting an excess of cardiac muscle cells and a defect of pharyngeal muscle cells in the mutant^3,23^. In the mouse, *Tbx1* appears to have a broader or earlier role, as its loss down regulates both cardiac and branchiomeric gene networks. Thus, rather than creating an imbalance between cardiac and branchiomeric lineages, the loss of *Tbx1* leads to a differentiation drift where a subset of cells expresses ectopic markers, such as the neuroectodermal protein OTX2 or the secreted extra cellular matrix protein periostin. The genetic labeling experiments showed that the up regulation of these two genes might be, at least in part, cell autonomous, and thereby suggesting that TBX1 suppresses an epithelial-like transcriptional program, in favor of a muscle fate. The finding of apoptotic activity in the PA-1 core mesoderm at a later (E10.5) developmental stage^24^ suggests that these inappropriately differentiating cells are eliminated by cell death.

We previously observed that in *Tbx1* mutants, the core mesoderm of PA-1 and the mesoderm in other segments of the pharyngeal apparatus was compacted and less prone to intermingle with other surrounding cell types^25^. This phenotype could be explained by the increased expression of adhesion molecules such as periostin, thereby conferring ectopic epithelial properties to the cells. Reduced expression of the essential extracellular matrix remodeling factor-encoding gene *Adamts9* ^26,27^ might complicate further the dynamics of mutant cells and affect regionalization of progenitors as well as cell-cell and cell-matrix signaling mechanisms^28^, as well as the diffusion of signals among different tissue types, such as mesoderm and neural crest-derived cells^29^. The hypothesis that the morphogenetic phenotype of *Tbx1* mutant mice is related to extra cellular matrix biology is supported by partial rescue of the heart phenotype using early embryonic treatment with the lysyl hydroxylase inhibitor Minoxidil^30^.

We propose that *Tbx1* is a priming factor for the CPM that modulates chromatin accessibility to genes within transcriptional modules involved in, a) development of cardiac and branchiomeric muscle, and b) regulation of morphogenetic processes. These modules appear to be distinct, as they are defined by different sets of networked genes. The mutant phenotype likely results from the perturbation of both differentiative and morphogenetic functions, and with a possible convergence downstream on ECM-cell or cell-cell interactions that affect cell positioning and movement, as well as the distribution of diffusible signals.

## MATERIALS AND METHODS

### Cell cultures, differentiation, and disaggregation

The mutant ES cell line 4D has been described previously^9^. This cell line, and its parental line ES-E14TG2a (ATCC CRL-1821), were cultured without feeder cells in GMEM (Sigma Cat# G5154) supplemented with 10^3^ U/ml ESGRO LIF (Millipore, Cat# ESG1107), 15% fetal bovine serum (ES Screened Fetal Bovine Serum, US Euroclone Cat# CHA30070L), 0.1 mM non-essential amino acids (Gibco, Cat# 11140-035), 0.1 mM 2-mercaptoethanol (Gibco, Cat# 31350-010), 0.1 mM L-glutamine (Gibco, Cat# 25030081), 0.1 mM Penicillin/Streptomycin (Gibco, Cat# 10378016), and 0.1 mM sodium pyruvate (Gibco, Cat# 11360-070). Cell passaging occurred every 2–3 days using 0.25% Trypsin-EDTA (1X) (Gibco, Cat# 25200056) as the dissociation buffer.

For differentiation, we followed a published protocol with minor modification ^8^. Briefly, cells were dissociated with Trypsin-EDTA and plated at a density of 100,000 cells/ml in serum-free differentiation media made of 75% Iscove’s modified Dulbecco’s media (Cellgro Cat# 15-016-CV) and 25% HAM F12 media (Cellgro #10-080-CV). The media were supplemented with N2 (GIBCO #17502048) and B27 (GIBCO #12587010), penicillin/streptomycin (GIBCO #10378016), 0.05% BSA (Invitrogen Cat#. P2489), L-glutamine (GIBCO #25030081), 5 mg/ml ascorbic acid (Sigma A4544), and 4.5 × 10-4M monothioglycerol (Sigma M-6145). After 48 hrs, the embryoid bodies (EB) were harvested and seeded in serum-free differentiation media supplemented with 1 ng/ml human Activin A (R&D Systems Cat#. 338-AC) and 1 ng/ml human BMP4 (R&D Systems Cat# 314-BP). Media were subsequently changed to serum-free differentiation media without additional growth factors after 2 days, and the EBs cultured until d6 or d8. EBs were left to settle to the bottom of 15ml tubes (∼2–5 minutes) before removing the media and the EBs were resuspended in DPBS without calcium and magnesium. EBs were disaggregated using the StemPro® Accutase® disaggregation kit (A1110501 Thermo Fisher) for 4-8min at room temperature. The disaggregation process was blocked by adding fresh pre-warmed complete medium. Cells were then washed multiple times DPBS without calcium and magnesium, resuspended in freezing media (20% FBS, 10% DMSO) and stored at −80 °C, prior to shipping in dry ice to the laboratory that performed the single cell multiomic assay (Berlin Institute of Health at Charité – Universitätsmedizin Berlin, Germany).

### Nuclei isolation and library generation

The cells were isolated and prepared for library generation following the 10x nuclei isolation protocol (CG000365, Rev C) for cell lines. Briefly, the cell pellets were thawed and cells were counted. After determining that more than 100,000 cells were present per sample the cells were lysed for 4 min. Afterwards, the isolated nuclei were washed and counted. Following the 10x Multiome protocol (CG000338, Rev E) the nuclei were diluted to load 8,000 nuclei per sample. The libraries were generated with the Multiome Kit (10x Genomics, PN-1000283) and the final libraries were sequenced on an Ilumina NovaSeq6000 (S4, paired-end). The sequencing data are deposited on the European Nucleotide Archive (Accession no.: PRJEB64050 [to be released]).

### Pre-processing and quality control

The sequencing data was aligned to the mouse genome (mm10/GRCm38) using cellranger-arc (v2.0.0) and afterwards aggregated. The data was filtered for quality control, subjected to unsupervised clustering, integrated and analyzed using the R packages Seurat ^31^ (v4.1.0) and Signac ^32^ (v1.6.0). During quality control, cells were filtered out, if they meet any one of these criteria: a) in RNA assay: < 200 genes OR < 1000 molecules OR > 30000 molecule OR > 10% mitochondrial genes; b) in ATAC assay: < 200 genes OR < 1000 molecules OR > 160000 OR > 10% mito genes. The dimensional reduction, clustering and integration was performed on both assays independently. Afterwards, a weighted nearest neighbors (WNN) graph was calculated combining the RNA and ATAC modalities.

### Multiomic single cell data handling and analyses

Given the integrated Seurat object and the results of clustering, we identified the top 25 markers for each cluster for annotating clusters with cell types using the function FindAllMarkers(min.pct= 0.2, only.pos = TRUE, logfc.threshold = 0.25, group.by= “wsnn_res.0.2”, test.use = “MAST”) from Seurat package (v. 4.4.0) ^31^. We assessed the significance of cell proportion within each cluster using the propeller function from the speckle package (v 1.2.0) ^33^.

For the CPM subset, we considered clusters ic1, ic4, ic8, and ic10. We recalculated the top 3000 variable genes using the “vst” method, scaled gene expression, and conducted dimension reduction with PCA employing 50 dimensions. Subsequently, we computed k-nearest neighbors with ndims=1:30, using the variable genes as features. We performed clustering analysis using the Louvain algorithm with a resolution parameter 0.30. We also re-identified the markers and assessed the significance of cell proportion with the method described above. We used the CPM population for the following further analysis.

### Multidimensional Scaling

We used a pseudo bulk approach to analyze the similarity between clusters in a lower-dimensional space using a MDS plot. First, we selected only the wild type genotype and we aggregated cell raw counts at the cluster-timepoint level summing all of the values in each group using muscat (v 1.14.0) ^34^. We then generated a multidimensional scaling (MDS) plot in which each point corresponds to a cluster-sample instance. The points are colored based on their cluster and shaped according to their group.

### Expression and enrichment analyses

We employed the Seurat approach to identify differentially expressed (DE) genes specific to each cluster for d6 and d8, comparing the genotypes. For this purpose, we utilized the FindMarkers function, choosing min.pct = 0.05 and logfc.threshold = 0.2. Similarly, we calculated differentially accessible regions (DARs) using the same function, with parameters specified for ATAC, i.e., min.pct = 0.05, test.use = ‘LR’, latent.vars = “nCount_ATAC”, where nCount_ATAC refers to the results of the ATAC assay. We visualized ATAC coverage using CoveragePlot function from Signac^32^.

### Motif analysis

We merged the list of DARs obtained from the analysis of all clusters and selected unique regions in case of repetition. We filtered out those overlapping with the blacklist regions (https://github.com/Boyle-Lab/ Blacklist/blob/master/lists/ mm10-blacklist.v2.bed.gz), and we ran HOMER (Hypergeometric Optimization of Motif EnRichment v. 4.11)^35^ on these regions with the parameters -size given using as background all the common scATAC peaks (“peakome”).

### Network and enrichment analysis

We employed high-dimensional weighted gene co-expression network analysis implemented in the hdWGCNA package (v.0.2.26)^14^ to construct a co-expression network based on single-cell data. Initially, we input genes expressed in at least 5% of the cells and utilized the MetacellsByGroups function to create the metacell gene expression matrix, with parameters set as k = 25 and min cells = 10. Subsequently, we applied the TestSoftPowers function to determine the optimal soft power and build the co-expression network using the ConstructNetwork function. Finally, we conducted a differential module eigengene (DME) analysis to identify modules that exhibit up or down-regulation in the KO vs WT groups of cells. We selected specific clusters and extracted the top 100 hub genes for modules of interest. We intersected top hub genes with genes annotated to scATAC peaks (extended +/− 15000 from the gene body). We calculated the coverage for each peak as follows: 1) Count fragments within the regions for different groups of cells (KO and WT for both days) using function RegionMatrix with the parameters upstream= 500, downstream = 500; 2) Normalize the matrix for cell number, 3) Convert the normalized matrix in a normalizedMatrix object using the function as.normalizedMatrix(k_upstream = 501, k_downstream = 500, k_target = 0, extend = 500, smooth = TRUE, keep = c(0, 0.95)) from ComplexHeatmap (v. 2.16.0)^36^ and visualize it in a heatmap. We evaluated the statistical significance between the compared groups using a one-sided paired Mann-Whitney test. Among these regions, we employed an L1-penalized logistic regression model to predict the probability of being an enhancer utilizing ATAC-seq data, as detailed in Aurigemma et al. 2024^37^. Briefly, we downloaded the enhancer coordinates (i.e., positive regions with response Y=1) from the VISTA ENHANCER Browser. Coordinates for non-enhancers (i.e., negative regions with response Y=0) were randomly generated, matching for cardinality and size those of enhancers. The feature matrix X was constructed with the available epigenetic tracks used by Aurigemma et al.^37^ (peak coordinates for histone modifications, p300 and CTCF transcription factors), assessing overlaps with both positive and negative regions. The regression model was trained using k-fold cross-validation. Subsequently, we classified a region as a potential enhancer by applying a threshold of 0.5 over the average estimation of the expected probability of being an enhancer. From this subset of potential enhancers, we further refined our selection by including only peaks with a LogFC (log-fold change) < −1.2 and a mean expression > 0.02. Subsequently, we employed Homer with the selected regions as input and setting size -given. For the background, we considered all peaks (peakome). We used the same method to create enrichment heatmaps for i) 255 TBX1 ChIP-seq peaks derived from mouse embryo tissue^12^ and ii) 2,388 ChIP-seq peaks derived from a cell culture system^11^.

We used Monocle3 (v1.3.7)^10,38^ package with default settings to conduct cell differentiation trajectory analysis on WT cell datasets.

For gene ontology searches we used the lists of hub genes from modules c1mod4, c1mod8, c3mod4, and the top 25 markers from c2 cluster, taken separately, as input for the gProfiler2 software (v0.2.3)^39^ to identify their functions and associated pathways. We set the background to include all expressed genes and used an FDR threshold < 0.01. For gene ontology search of DAR-associated genes, we merged all DARs identified for each cluster, removing repetitions. We annotated them to genes using the makeTxDbFromEnsembl function from ChIPseeker (v1.38.0)^40^ by associating to each peak/region the nearest gene, setting the TSS region [−3000, 3000] and using the release 102 from the mus musculus Ensembl database. We then used the annotated genes as input for gProfiler2, setting the background to genes annotated to peaks with a total number of fragments > 500 in the cells of the clusters of interest.

For RNA velocity-based tests, we used loom files containing spliced and unspliced counts, derived from bam files, as input data for Dynamo (v. 0.95.2)^21^ to infer cell trajectories. Using the adata.concatenate function, we merged the loom files for KO and WT cells separately, resulting in two genotype-specific independent analyses. Subsequently, we employed the “stochastic” model within Dynamo to calculate RNA velocity for each cell for both genotypes. By reducing the dimensionality of the high-dimensional velocity vectors and reconstructing vector fields, we gained insights into the topology of the CPM vector field. To characterize comprehensively the regulatory mechanism, we also computed cell acceleration, which represents the derivative of the velocity vector. Positive acceleration indicates cell speeding up, while negative values indicate deceleration. We visualized the acceleration of selected marker genes in UMAP to compare them visually across both conditions.

### Immunofluorescence

The embryos were fixed in 4% PFA, embedded in paraffin, and sectioned at a thickness of 10 µm. The sections were rehydrated, and antigen retrieval was performed in 10 mM sodium citrate pH 6.0. Blocking was carried out using 0.5% milk, 10% FBS, and 1% BSA (bovine serum albumin) in H_2_O (Blocking solution). Incubation with the primary antibody in blocking solution was carried out overnight at room temperature, and signal detection was performed using secondary antibodies conjugated with different fluorochromes (1:500). We used the following primary antibodies: Anti-GFP #ab13970 (Abcam) 1:1000; Anti-Periostin/OSF2 #NBP1-30042 (Novusbio) 1:100; Anti-Otx2 #AF1979 (R&D Systems) 1:100; The secondary antibodies were: Donkey anti-Goat IgG (H+L), Alexa Fluor™ 488 #A11055; Donkey anti-Chicken IgY (H+L), Alexa Fluor™ 594 #A78951; Donkey anti-Rabbit IgG (H+L) Alexa Fluor™ 594 #A21207. Imaging was carried out using a Leica Thunder Imaging System (Leica Microsystems) equipped with a Leica DFC9000 GTC Camera, lumencor fluorescence LED light source, and the software Leica Application Suite X 3.7.4.23463.

### RNAscope *in situ* hybridization

Embryos were fixed in 4% PFA, dehydrated with increasing concentrations of methanol, and stored at −20°C. For the hybridization we followed the protocols suggested by the provider of the RNAscope reagents (RNAscope™, ACD-Biotechne); briefly, whole embryos were rehydrated and permeabilized for 20 minutes with Protease III, followed by overnight incubation at 40°C with the probe solution (see list below) and the signal was developed with TSA-FITC (Akoya Bioscience NEL741001KT, 1:500) and TSA-CY3 (Akoya Bioscience NEL745001KT, 1:2000). The embryos were cryoprotected with increasing concentration of sucrose (10%, 20%, and 30% sucrose/PBS 1X) at 4°C, OCT/sucrose 30% (50:50), embedded in OCT, sectioned at a thickness of 10 µm, and imaged using the Leica Thunder Imaging System described above. We used the following commercial probes provided by RNAscope™: Probe-Mm-Postn Cat No. 418581; Probe-Mm-Grid2 Cat No. 512701; Probe-Mm-Tbx1-C2 Cat No. 481911-C2; Probe-Mm-Adamts9 Cat No. 400441; Probe-Mm-Tshz2 Cat No. 431061. Imaging was carried out as for immunofluorescence experiments.

### Standard RNA *in situ* hybridization

For *in situ* hybridization with mouse *Otx2* (a kind gift from the laboratory of Dr. P. Chambon), the probe was reverse-transcribed and labeled with Digoxigenin using the digoxigenin RNA labeling kit (Merck, Cat. 11277073910). Incubation with the probe was performed overnight at 60°C on whole mount embryos, blocking with 10% Sheep serum in 1x PBS, and incubation with Anti-Digoxigenin-AP (Merck Cat. 11093274910) diluted 1:2000 in blocking solution overnight at 4°C. The embryos were then cryoprotected with increasing concentrations of sucrose (10%, 20%, and 30% sucrose/PBS 1X) at 4°C, OCT/sucrose 30% (50:50), and embedded in OCT. Immunostained embryos were then cryosectioned at a thickness of 10 µm for observation and imaging using a Nikon Eclipse Ni microscope equipped with a Nikon DS-RI1 Camera and Nis Elements AR 4.20.00 software.

## AUTHOR CONTRIBUTIONS

OL: Bioinformatic strategies and analyses; MB: mouse breeding and handling, immunofluorescence, RNAscope, RF: cell culture and differentiation; IA: gene editing; EI: funding recruitment, manuscript editing; CA: bioinformatic analysis supervision, manuscript editing; AB: project conceptualization, funding recruitment, manuscript writing; JL: Sample Prep, raw data handling, QC, integration, data handling, bioinformatic support; KJ: Sample Prep, library prep; SL: bioinformatic support and input; FT: Data handling, bioinformatic support.

## ACKNOWLEDGEMENTS

We thank Giuseppina Divisato at the Microscopy core of the Department of Molecular Medicine and Medical Biotechnology, Univ. Federico II; Lucia Mele for technical help with mouse handling; Varsha Poondi Krishnan for initial help with bioinformatic data handling; the microscopy core of the IGB-CNR. This project has received funding from the European Union’s Horizon 2020 Research and Innovation Programme under grant agreement No 824110; the German Ministry of Education and Research 01ZZ2001; the Telethon Foundation GMR22T1012, and from the Italian Ministry of University and Research PRIN 2022XFE7M2.

## SUPPLEMENTARY TABLES

**Supplementary Table 1**

*Top 25 marker genes of clusters of the full dataset and of the CPM subset*.

Sheet 1: top 25 markers of clusters of full dataset. Sheet 2: top 25 markers of clusters of the CPM subset.

**Supplementary Table 2**

*Differentially accessible regions (DARs) identified in the CPM subset and annotated with the neighboring gene*.

Sheet 1: Full list of 185 unique DARs identified in the CPM subset Sheets 2-7: DARs identified in the indicated clusters.

**Supplementary Table 3**

*Search results of motif enrichment analysis using the full set of DARs of the CPM subset (n=185). We used Homer and reported only the significantly enriched motifs*.

**Supplementary Table 4**

*Gene Ontology (GO) search results*.

Sheet 1: GO search of genes mapped to the DARs.

Sheets 2-4: GO search results of top Hub genes of the transcriptional modules indicated.

Sheet 5: GO search of marker genes of cluster c2 of the CPM subset.

**Supplementary Table 5**

*Differentially expressed genes in clusters of the CPM subset*.

**Supplementary Tab. 6**

*Hub genes resulting from hdWCGNA analysis of co-expressed genes in the CPM subset divided by clusters. The Log2FC and adjusted P values refer to differences between ko and wt cells*.

**Supplementary Table 7**

*Putative enhancers mapped to hub genes of transcription modules c1mod4, c1mod8, and c3mod4. Prediction scores and scATAC-seq coverage changes in ko vs. wt cells are indicated*.

## SUPPLEMENTARY FIGURES

**Supplementary Figure 1**

*Major lineage composition and effects of differentiation time and genotype on the population*

a) Markers of the FHF (*Hand1*, blue) and SHF (*Tbx1*, red).

b) Left, clusters expressing SHF markers (in blue), and FHF markers (in red); right, names of clusters.

c) The expression of *Tbx1* has a similar distribution at the different differentiation time points.

d) Left, the effect of genotype (WT vs KO) is visually limited; right, the effect of differentiation time is more visible, especially on clusters ic1, ic2, ic4.

**Supplementary Figure 2**

*The cell differentiation model expresses the major markers of CPM differentiation but does not express posterior second heart field (pSHF) marker genes*.

a-e) The CPM dataset includes the most typical cell types in this population: cardiac progenitors (a), cardiomyocytes (b), smooth muscle cells (c) branchiomeric muscle progenitors (d), and a small population of branchiomeric muscle cells (e),

f-g) *Hoxb1* and *Tbx5*, markers of pSHF, are not expressed in the CPM dataset. h) *Tbx5* is expressed in the full dataset but it is restricted to the FHF domain.

**Supplementary Figure 3**

*Interrelations between clusters of the CPM subset*.

a) Multidimensional scaling measures the Euclidean distance between the transcriptomes of different clusters, note the vicinity between clusters c1-c3 and c0-c2 (circled).

b) Pseudotime analysis using Monocle3 also identifies connections between c0 (d6) and c2 (d8) as well as between c1 (d6) and c3 (d8) (arrows).

**Supplementary Figure 4**

*Examples of scATAC coverage of selected DARs of the CPM subset*.

**Supplementary Figure 5**

*Heatmaps of gene expression of hub genes of the transcriptional modules c1mod4, c1mod8, and c3mod4*.

**Supplementary Figure 6**

*Transcriptional modules in the CPM subset: late time point*.

a) Connections among hub genes of module c3mod4, a d8 module. The inner circle includes the most connected genes.

b) Chromatin status of hub genes of c3mod4 responds significantly to loss of the *Tbx1* gene at promoters (left panels), and over the entire gene body (all ATAC peaks, right panels).

c) Enriched binding motifs identified in all the scATAC peaks (n=711) mapped to the c3mod4 hub genes in the CPM subset.

d) Enriched binding motifs identified in the scATAC peaks that respond to loss of *Tbx1*(n=131) and mapped to the c3mod4 hub genes in the CPM subset.

**Supplementary Figure 7**

*The genes encoding the myogenic regulatory factors Msc and Myf5 are not detectable in mutant cells of the CPM subset*.

**Supplementary Figure 8**

*Tshz2 gene expression in E9.5 mouse embryos assayed by RNAscope*.

a) 2-color RNAscope on a sagittal section reveals the substantial overlap of expression between *Tshz2* and *Tbx1*.

b) RNAscope on sagittal sections of mouse E9.5 WT and *Tbx1^−/−^* embryos. Note the reduced expression of *Tshz2* in the core mesoderm of the first pharyngeal arch of the mutant. Overall, the expression pattern of the gene is rearranged in the mutant, at least in part due to anatomical differences. Scale bars 100µm.

**Supplementary Figure 9**

*Genes encoding the CPM transcription factors Tcf21, Eya1, and Eya4 are affected by loss of Tbx1. ATAC coverage and RNA expression*.

a-a’) Regions of reduced accessibility in the vicinity of genes *Tcf21* and *Six1*.

b-b’) Reduced gene expression but apparently no neighboring chromatin changes for genes *Eya1* and *Eya4* encoding co-factors of SIX transcription factors.

**Supplementary Figure 10**

*The neuroectodermal gene Otx2 is expanded in the mutant CPM subset*.

a) *Otx2* gene expression is expanded in mutant cells of the CPM subset (arrows).

b) ATAC coverage shows increased opening of the promoter region of the *Otx2* gene in mutant c3 cells.

c) Sagittal sections of E8.5 (16 somites) and E9.5 (26 somites) mouse embryos with the indicated genotype, stained by immunofluorescence with an anti-OTX2 antibody. Normally, OTX2 is detectable in the nervous system and in the ectoderm of the first and second pharyngeal arches (PA-1 and PA-II). In the mutant embryos, the protein is also clearly detectable in the head mesenchyme and mesodermal core of PA-I (PA-II does not develop in the mutant). Scale bars are 100 µm.

**Supplementary Figure 11**

*Otx2 gene expression is up regulated in the core mesoderm of the first pharyngeal arch of Tbx1 mutant embryos*.

*In situ* hybridization using an *Otx2* probe on sagittal sections of E9.5 mouse embryos, the arrows indicate the first pharyngeal arch.

Scale bar is 100 µm.

**Supplementary Figure 12**

*Genetic labelling of Tbx1-expressing cells and their descendants reveals partial overlap with ectopic OTX2-expressing cells in homozygous Tbx1 mutant embryos*.

Transverse sections of E9.5 mouse embryos stained with two-color immunofluorescence anti-GFP (genetic label of *Tbx1*-expressing cells, in red) and anti-OTX2 (in green): a) control (heterozygous) embryo, merged image (left panel) and split channels green and red (right panels); b) Homozygous mutant embryo, note the region of up regulation in the first pharyngeal arch and head mesenchyme (arrows).

Scale bars are 100 µm.

**Supplementary Figure 13**

*Genetic labelling of Tbx1-expressing cells and their descendants reveals partial overlap with ectopic OTX2-expressing cells in homozygous mutant embryos (continued)*.

Sagittal sections of E9.5 mouse embryos stained with two-color immunofluorescence anti-GFP (genetic label of *Tbx1*-expressing cells, in red) and anti-OTX2 (in green): a) control (heterozygous) embryo, merged image (left panel) and split channels green and red (right panels); b) Homozygous mutant embryo, note the region of up regulation in the first pharyngeal arch (arrow).

Scale bars are 100 µm.

