## Supplementary Figures 1 to 13 for "*Tbx1* stabilizes differentiation of the cardiopharyngeal mesoderm and drives morphogenesis in the pharyngeal apparatus"

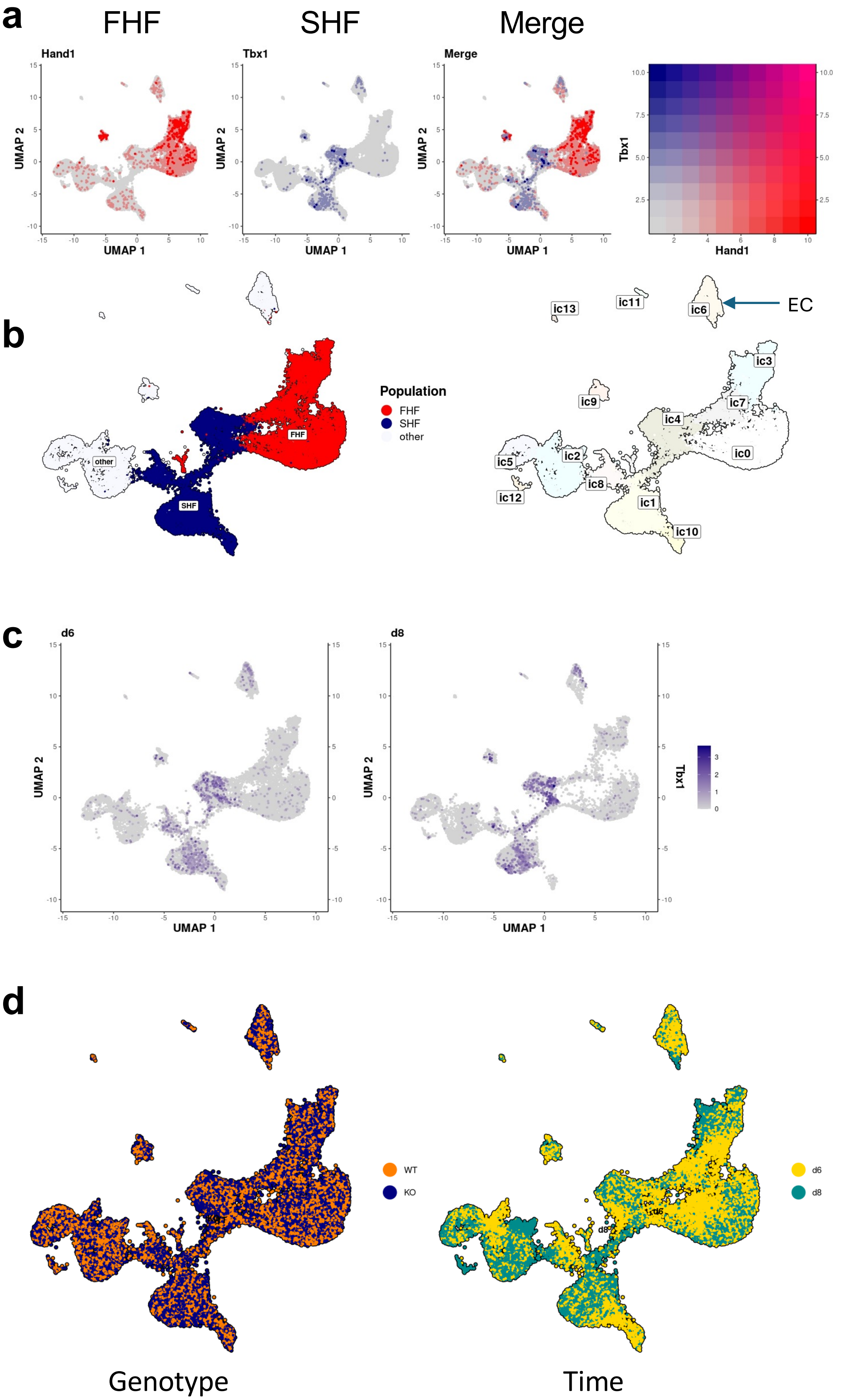


#### Supplementary Figure 1

*Major lineage composition and effects of differentiation time and genotype on the population*

a) Markers of the FHF (*Hand1*, blue) and SHF (*Tbx1*, red).

b) Left, clusters expressing SHF markers (in blue), and FHF markers (in red); right, names of clusters.

c) The expression of *Tbx1* has a similar distribution at the different differentiation time points.

d) Left, the effect of genotype (WT vs KO) is visually limited; right, the effect of differentiation time is more visible, especially on clusters ic1, ic2, ic4.

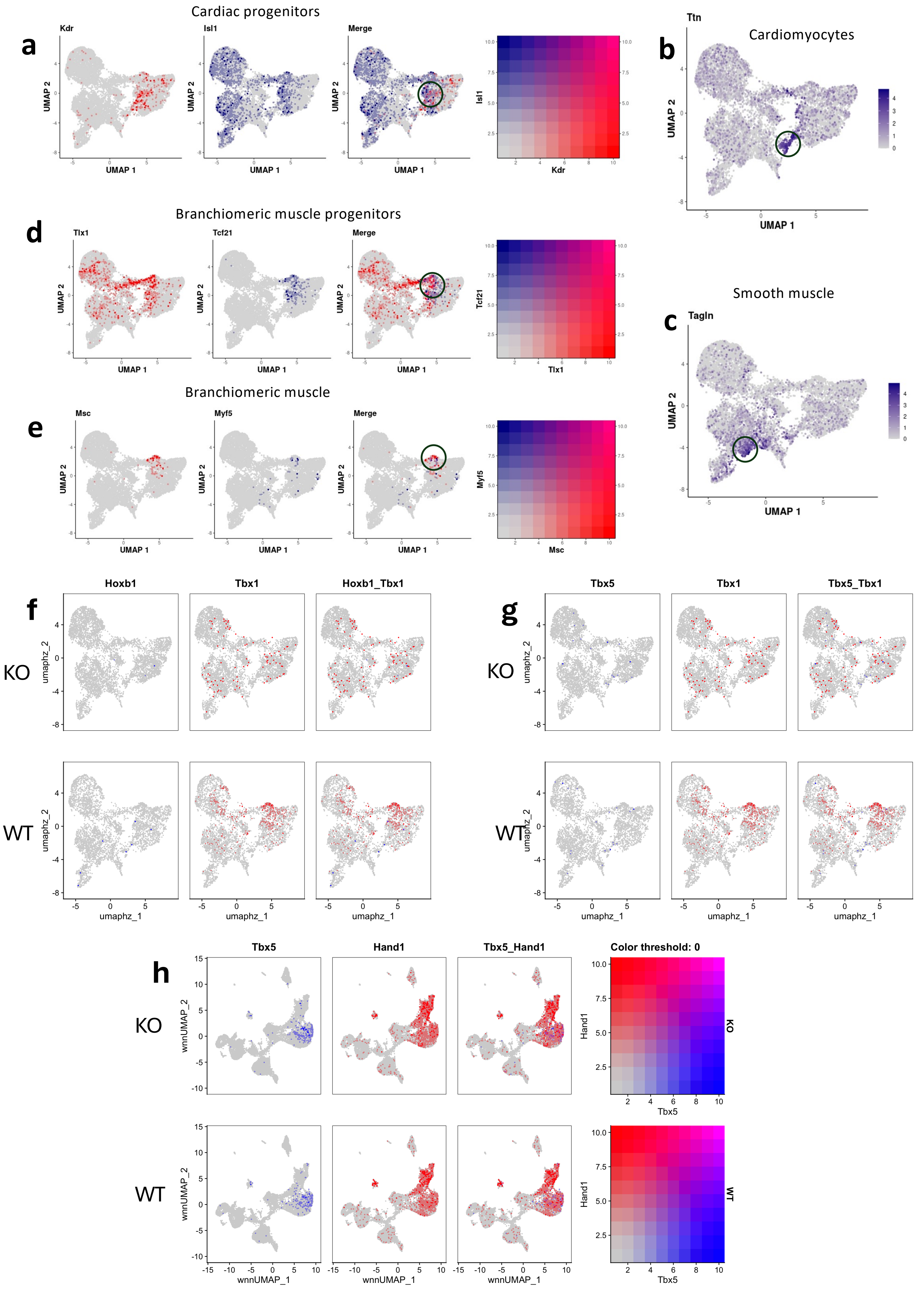


### Supplementary Figure 2

#### Legend to supplementary Figure 2

*The cell differentiation model expresses the major markers of CPM differentiation but does not express posterior second heart field (pSHF) marker genes.*
a-e) The CPM dataset includes the most typical cell types in this population: cardiac progenitors (a), cardiomyocytes (b), smooth muscle cells (c) branchiomeric muscle progenitors (d), and a small population of branchiomeric muscle cells (e),
f-g) *Hoxb1* and *Tbx5*, markers of pSHF, are not expressed in the CPM dataset. h) *Tbx5* is expressed in the full dataset but it is restricted to the FHF domain.

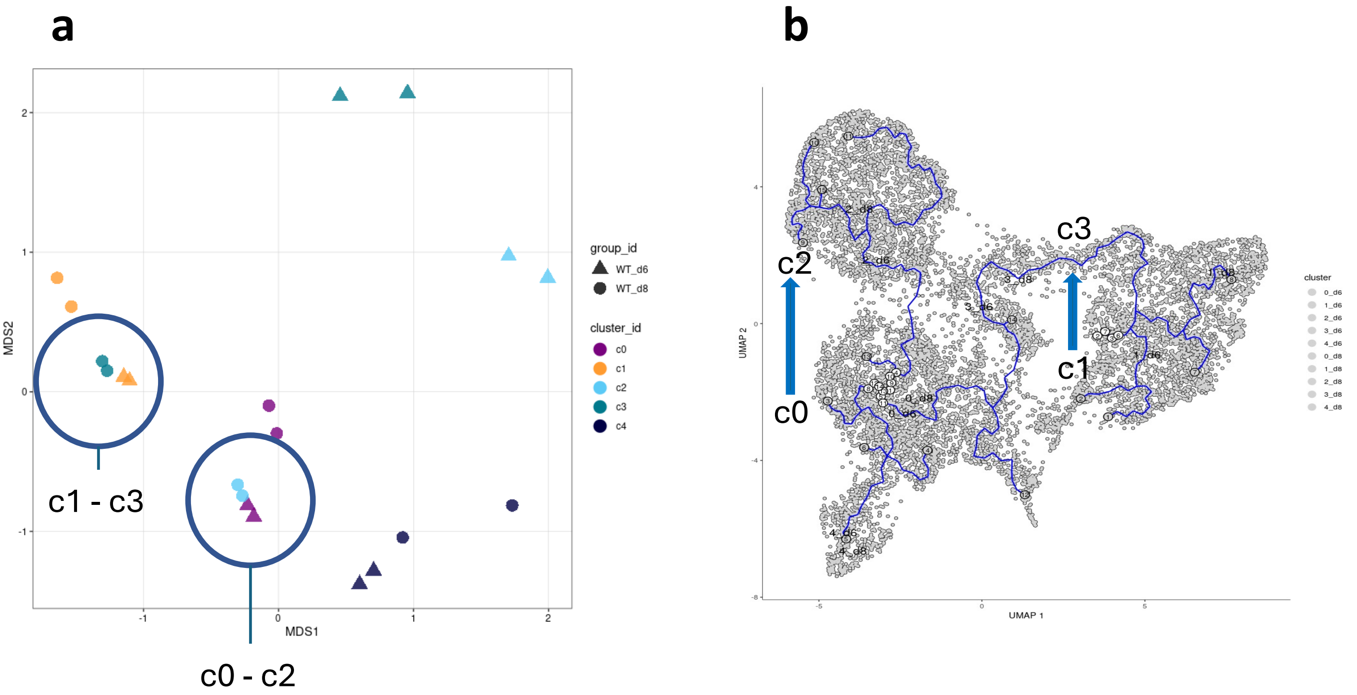


### Supplementary Figure 3

*Interrelations between clusters of the CPM subset.*

a) Multidimensional scaling measures the Euclidean distance between the transcriptomes of different clusters, note the vicinity between clusters c1-c3 and c0-c2 (circled).

b) Pseudotime analysis using Monocle3 also identifies connections between c0 (d6) and c2 (d8) as well as between c1 (d6) and c3 (d8) (arrows).

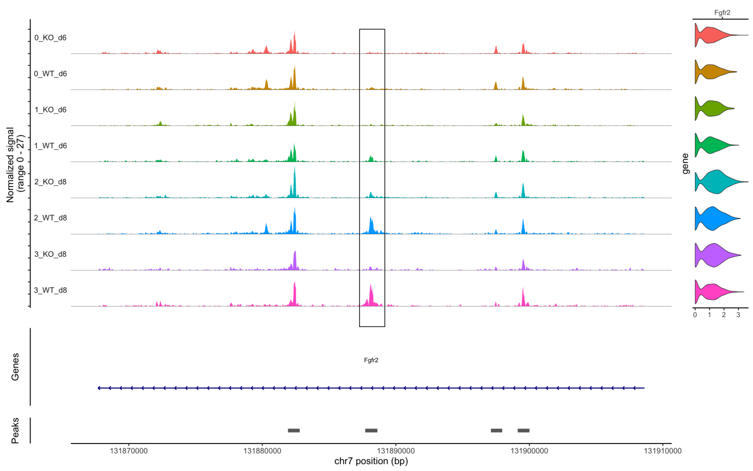


chr7:131887706-131888623

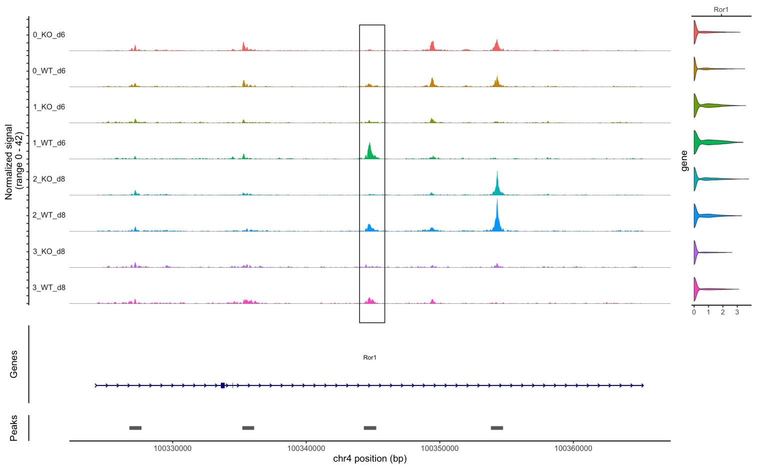


chr4:100344316-100345244

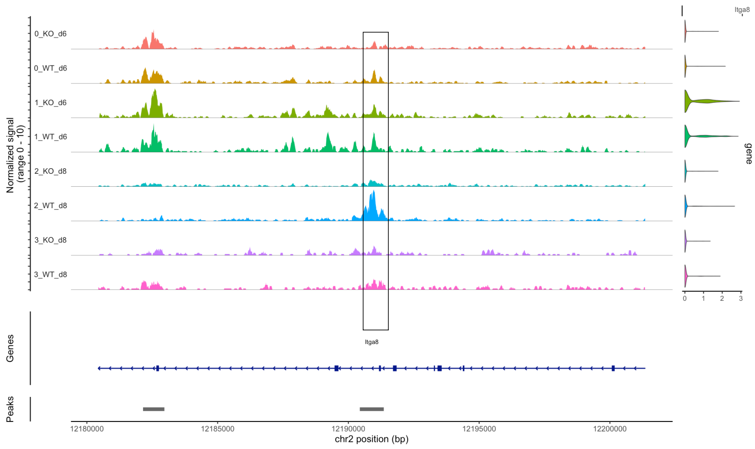


chr2:12190437-12191354

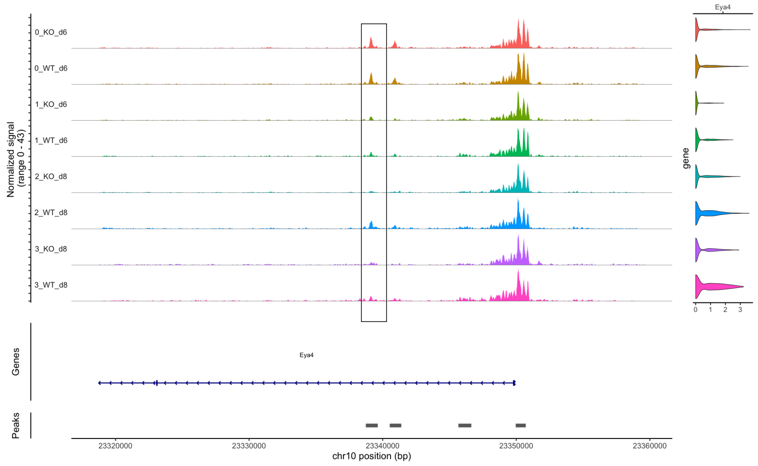


chr10:23338738-23339634

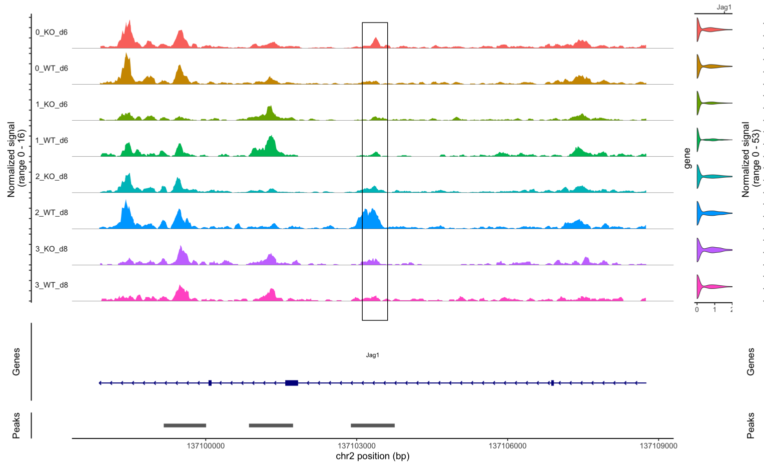


chr2:137102885-137103756

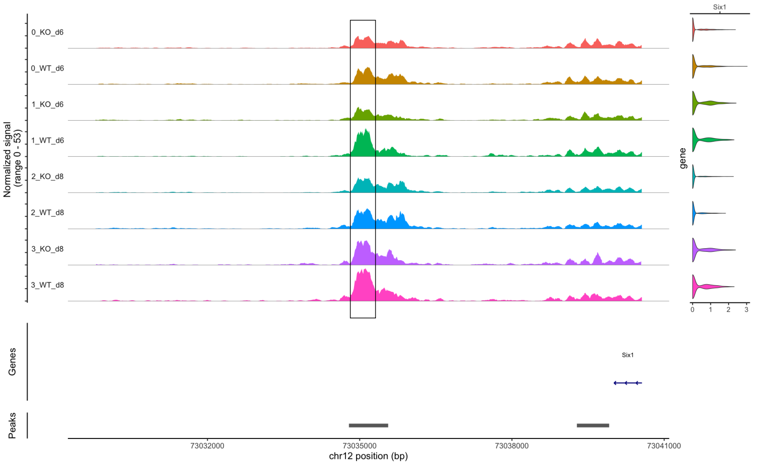


chr12:73034788-73035561

### Supplementary Figure 4

*Examples of scATAC coverage of selected DARs of the CPM subset.*

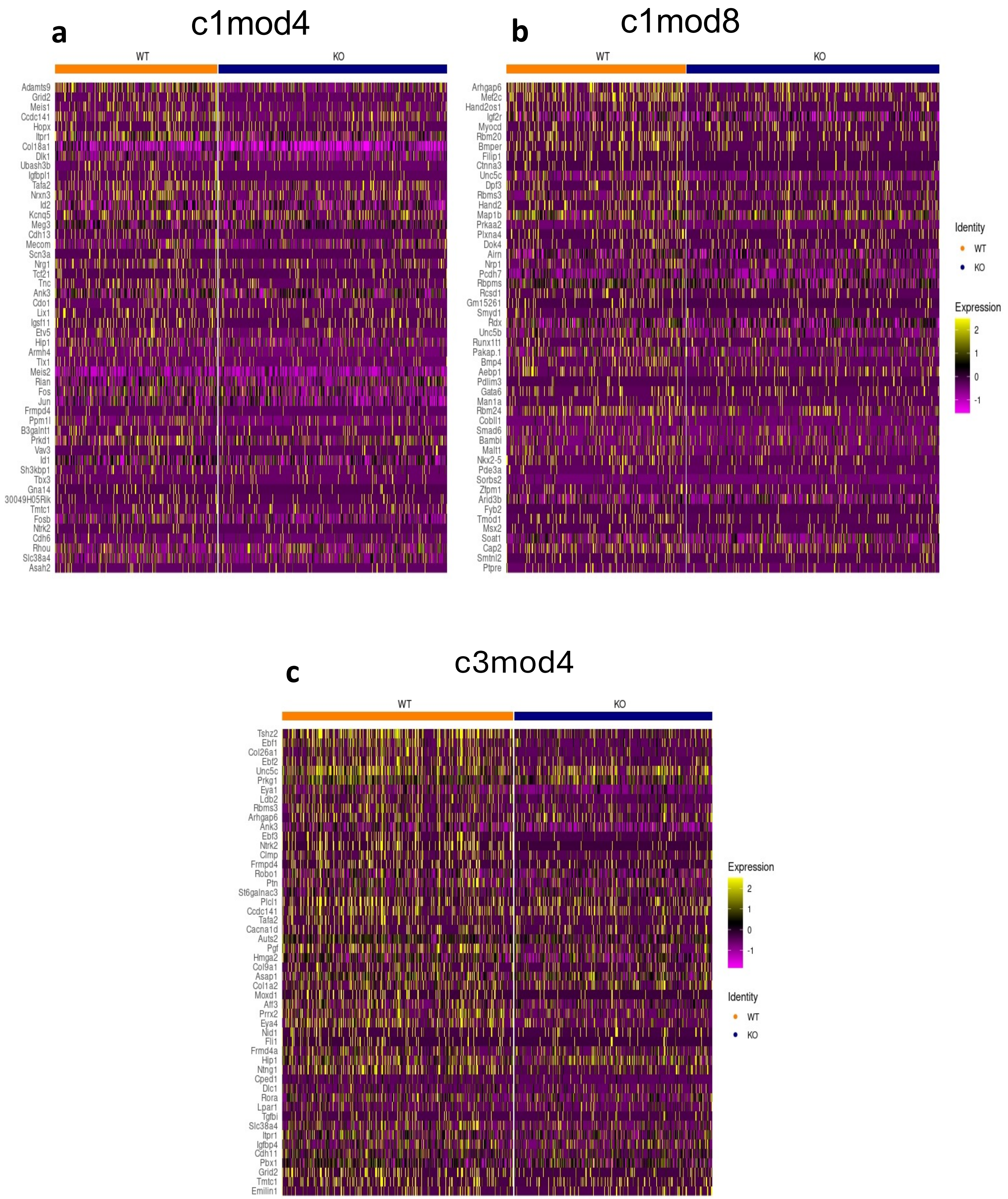


**Supplementary Figure 5**

*Heatmaps of gene expression of hub genes of the transcriptional modules c1mod4, c1mod8, and c3mod4.*

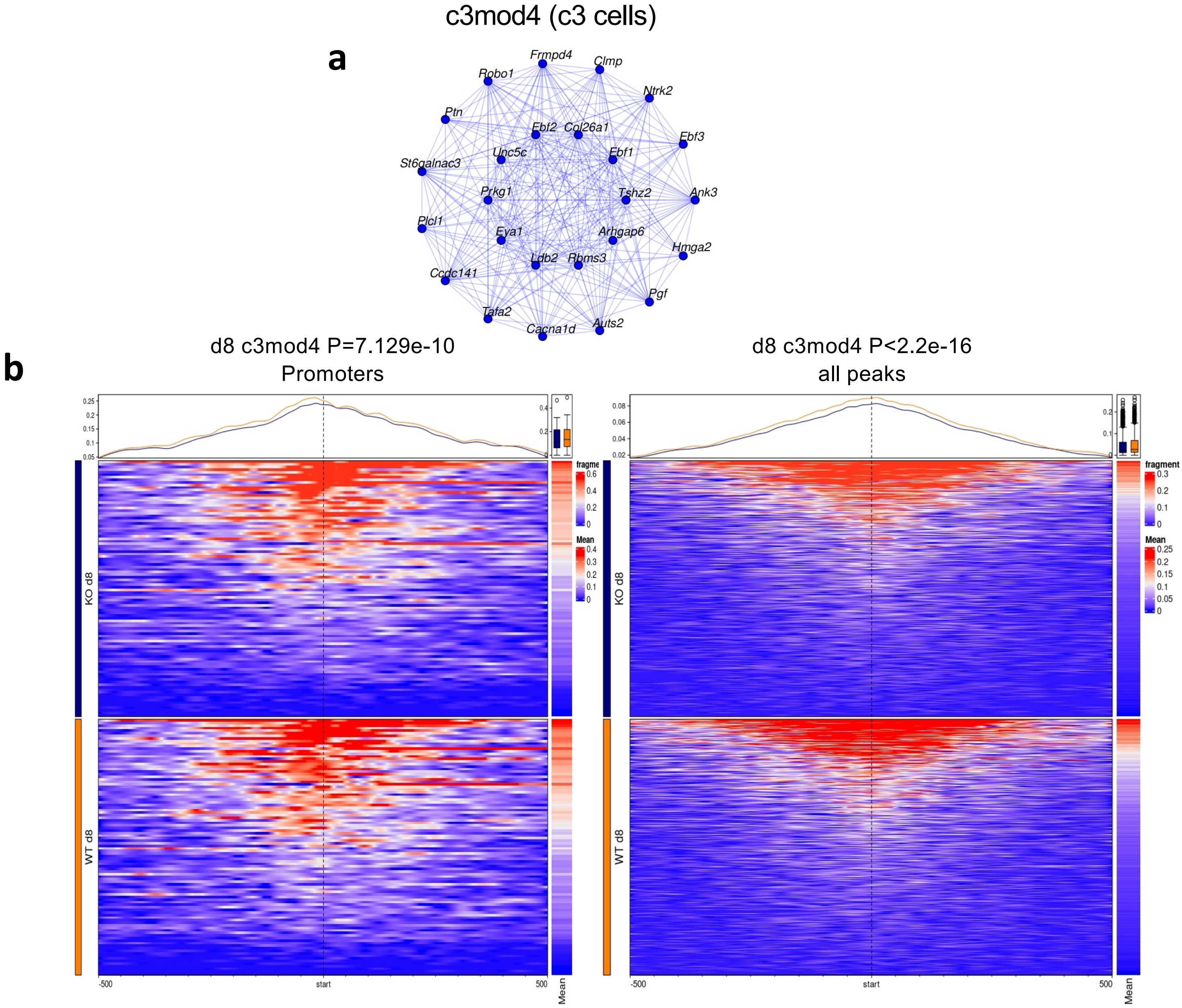


c

Total Target Sequences = 711, Total Background Sequences = 129343

| Rank | Motif | Name | P-value | log P-pvalue | q-value (Benjamini) | # Target Sequences with Motif | % of Targets Sequences with Motif | # Background Sequences with Motif | % of Background Sequences with Motif |
|---|---|---|---|---|---|---|---|---|---|
| 1 | | Pitx1:Ebox(Homeobox,bHLH)/Hindlimb-Pitx1-ChIP-Seq(GSE41591)/Homer | 1e-7 | -1.681e+01 | 0.0000 | 87.0 | 12.24% | 8627.2 | 6.67% |
| 2 | | EWS:ERG-fusion(ETS)/CADO_ES1-EWS:ERG-ChIP-Seq(SRA014231)/Homer | 1e-4 | -9.297e+00 | 0.0202 | 243.0 | 34.18% | 35831.9 | 27.70% |
| 3 | | NFAT:AP1(RHD,bZIP)/Jurkat-NFATC1-ChIP-Seq(Jolma_et_al.)/Homer | 1e-3 | -8.394e+00 | 0.0332 | 68.0 | 9.56% | 7918.8 | 6.12% |

d

Total Target Sequences = 131, Total Background Sequences = 128678

| Rank | Motif | Name | P-value | log P-pvalue | q-value (Benjamini) | # Target Sequences with Motif | % of Targets Sequences with Motif | # Background Sequences with Motif | % of Background Sequences with Motif |
|---|---|---|---|---|---|---|---|---|---|
| 1 | | Six1(Homeobox)/Myoblast-Six1-ChIP-Chip(GSE20150)/Homer | 1e-6 | -1.485e+01 | 0.0002 | 34.0 | 25.95% | 13331.0 | 10.36% |
| 2 | | Six2(Homeobox)/NephronProgenitor-Six2-ChIP-Seq(GSE39837)/Homer | 1e-4 | -9.632e+00 | 0.0144 | 68.0 | 51.91% | 45301.5 | 35.20% |
| 3 | | Pitx1:Ebox(Homeobox,bHLH)/Hindlimb-Pitx1-ChIP-Seq(GSE41591)/Homer | 1e-3 | -8.903e+00 | 0.0199 | 21.0 | 16.03% | 8470.7 | 6.58% |

#### Supplementary Figure 6

*Transcriptional modules in the CPM subset: late time point.*

a) Connections among hub genes of module c3mod4, a d8 module. The inner circle includes the most connected genes. b) Chromatin status of hub genes of c3mod4 responds significantly to loss of the *Tbx1* gene at promoters (left panels), and over the entire gene body (all ATAC peaks, right panels).

c) Enriched binding motifs identified in all the scATAC peaks (n=711) mapped to the c3mod4 hub genes in the CPM subset. d) Enriched binding motifs identified in the scATAC peaks that respond to loss of *Tbx1*(n=131) and mapped to the c3mod4 hub genes in the CPM subset.

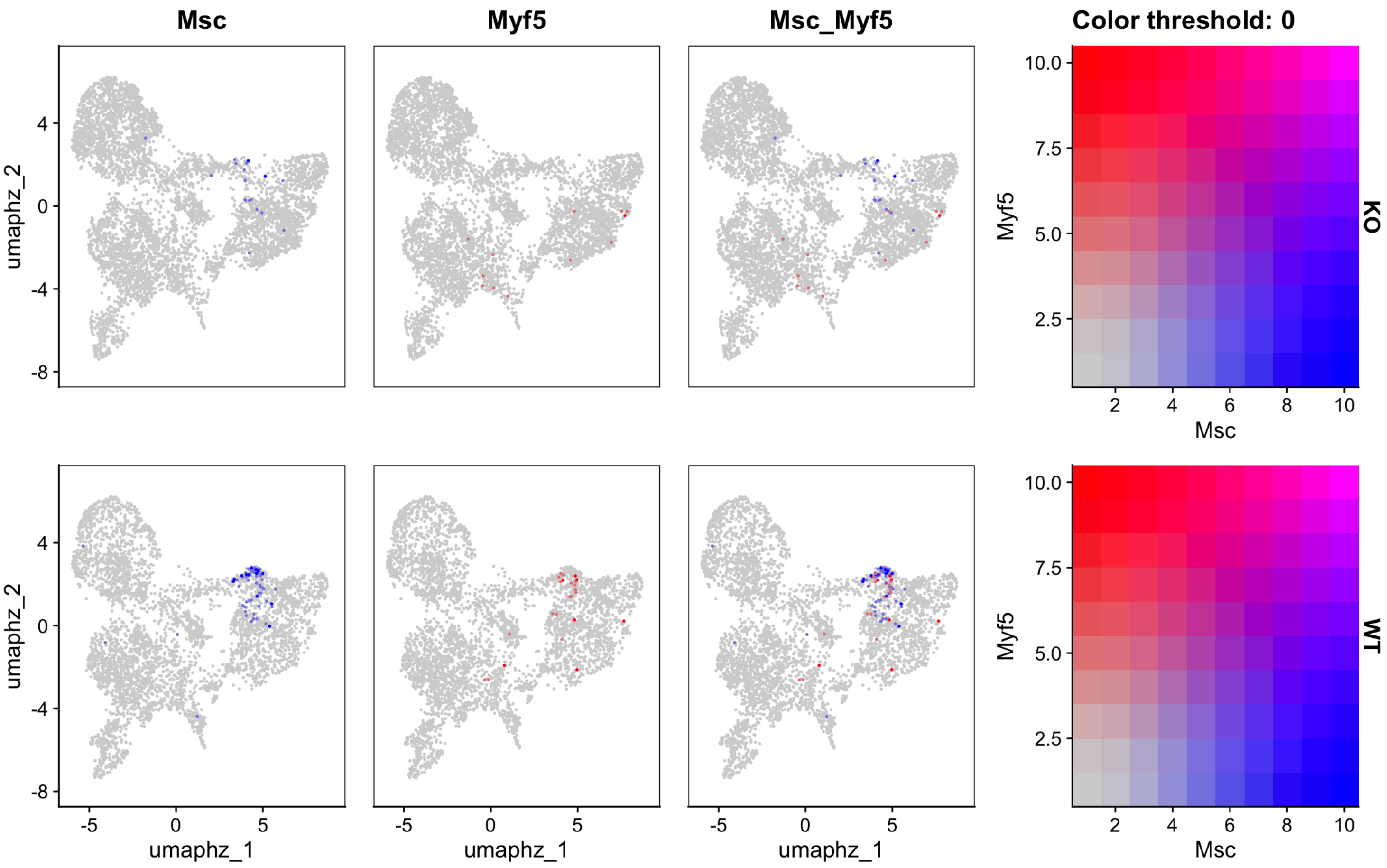


#### Supplementary Figure 7

*The genes encoding the myogenic regulatory factors Msc and Myf5 are not detectable in mutant cells of the CPM subset.*

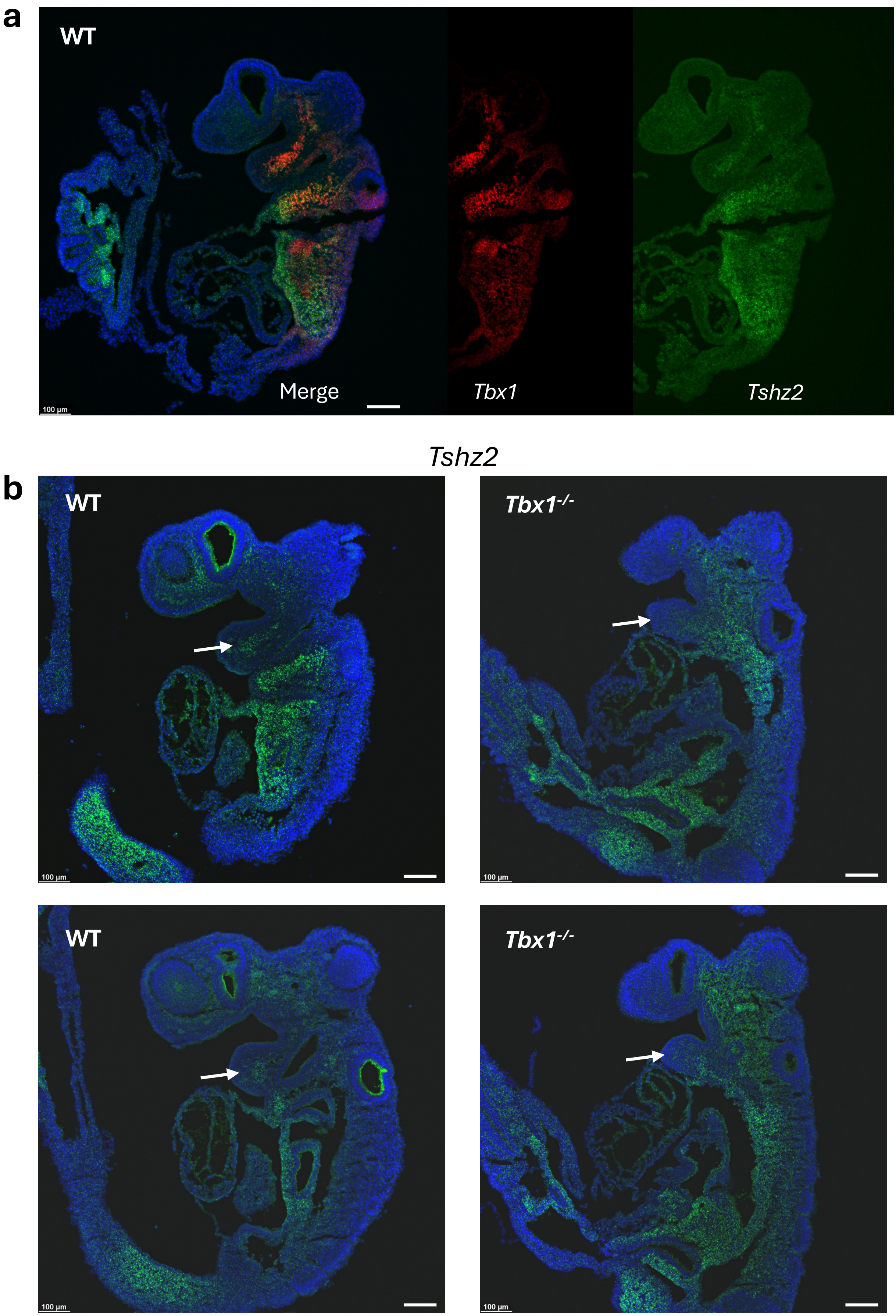


#### Supplementary Figure 8

*Tshz2 gene expression in E9.5 mouse embryos assayed by RNAscope.*

a) 2-color RNAscope on a sagittal section, there is a substantial overlap of expression patterns between the *Tshz2* and *Tbx1*.

b) RNAscope on sagittal sections of mouse E9.5 WT and Tbx1-/- embryos. Note the reduced expression of *Tshz2* in the core mesoderm of the first pharyngeal arch of the mutant. Overall, the expression pattern of the gene is rearranged in the mutant, at least in part due t anatomical differences. Scale bars 100µm.

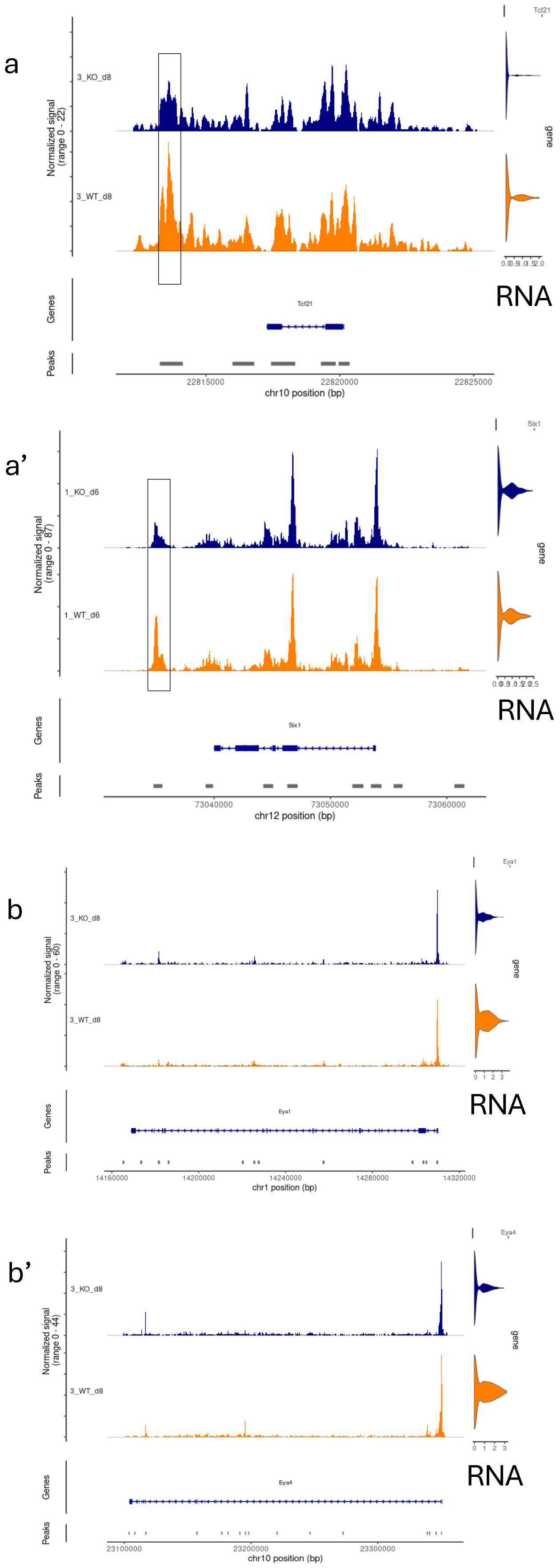


### Supplementary Figure 9

*Genes encoding the CPM transcription factors Tcf21, Eya1, and Eya4 are affected by loss of Tbx1. ATAC coverage and RNA expression.*

a-a') Regions of reduced accessibility in the vicinity of genes *Tcf21* and *Six1*.

b-b') Reduced gene expression but apparently no neighboring chromatin changes for gene *Eya1* and *Eya4* encoding co-factors of SIX transcription factors.

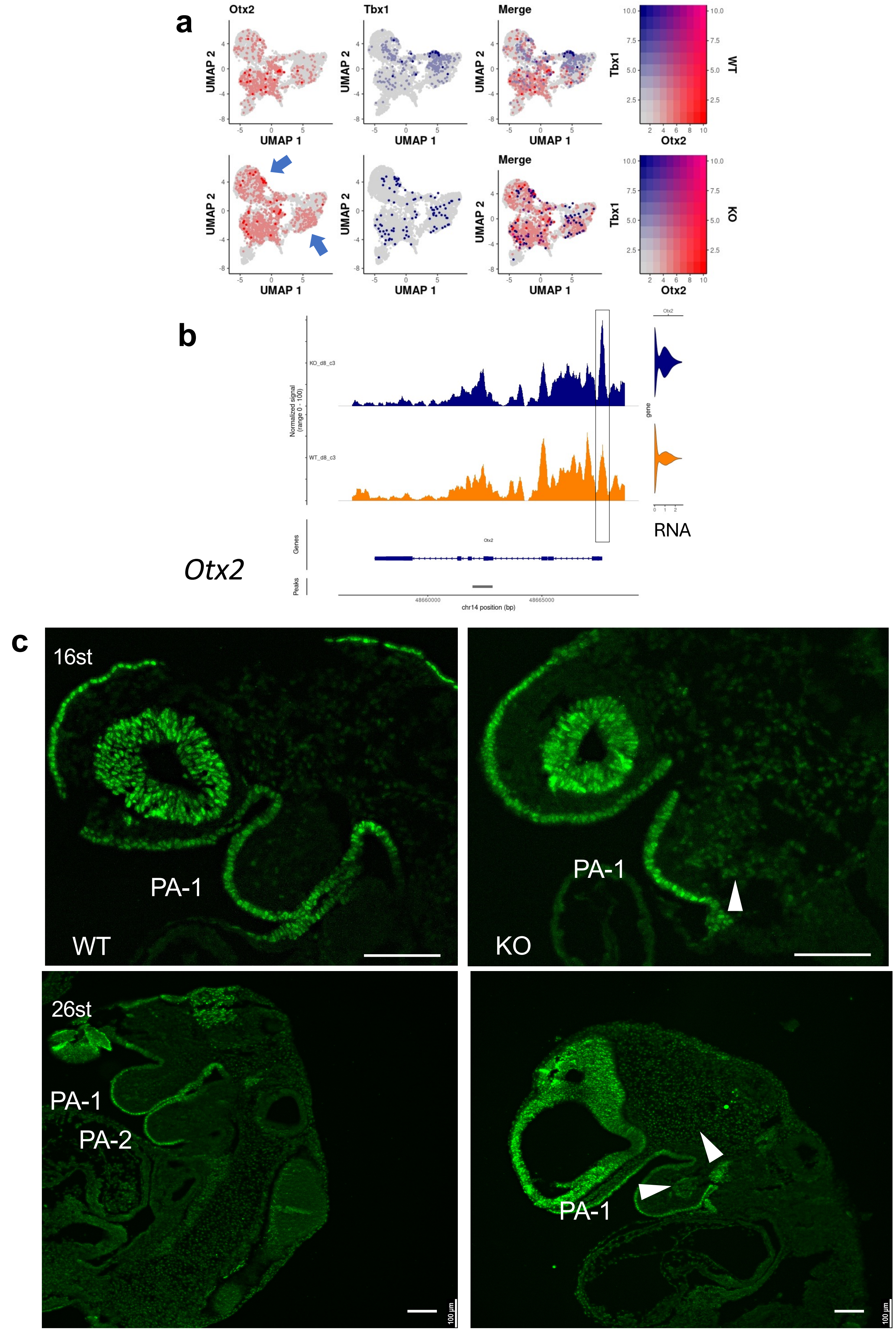


**Supplementary Figure 10**

#### Legend to supplementary Figure 10

*The neuroectodermal gene Otx2 is expanded in the mutant CPM subset.*

a) *Otx2* gene expression is expanded in mutant cells of the CPM subset (arrows).

b) ATAC coverage shows increased opening of the promoter region of the *Otx2* gene in mutant c3 cells.

c) Sagittal sections of E8.5 (16 somites) and E9.5 (26 somites) mouse embryos with the indicated genotype, stained by immunofluorescence with an anti-OTX2 antibody. Normally, OTX2 is detectable in the nervous system and in the ectoderm of the first and second pharyngeal arches (PA-1 and PA-II). In the mutant embryos, the protein is also clearly detectable in the head mesenchyme and mesodermal core of PA-I (PA-II does not develop in the mutant). Scale bars are 100 µm.

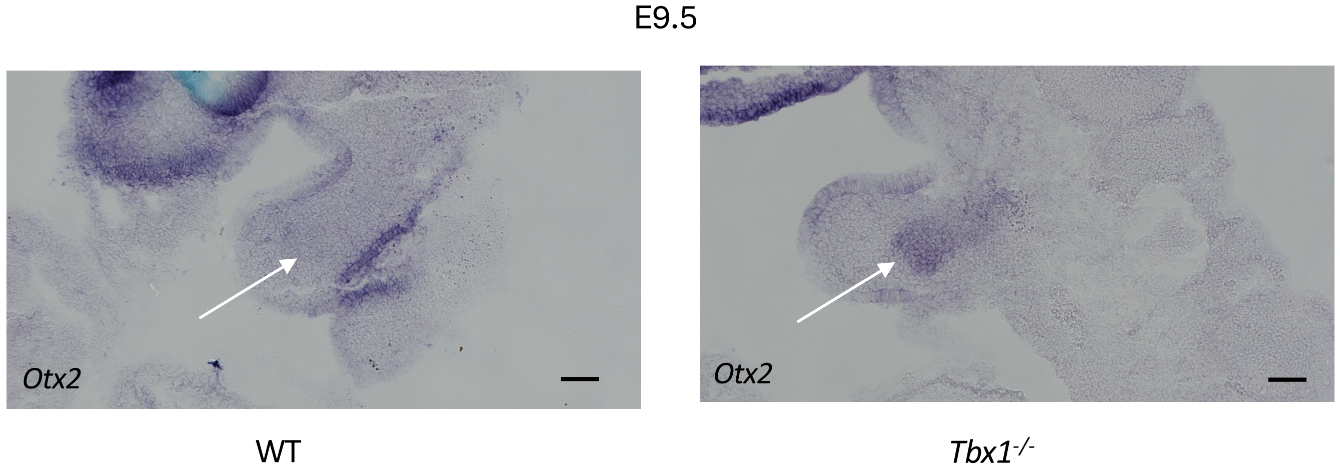


#### Supplementary Figure 11

*Otx2 gene expression is up regulated in the core mesoderm of the first pharyngeal arch of Tbx1 mutant embryos.*

In situ hybridization using an *Otx2* probe on sagittal sections of E9.5 mouse embryos, the arrows indicate the first pharyngeal arch.

Scale bar is 100 μm.

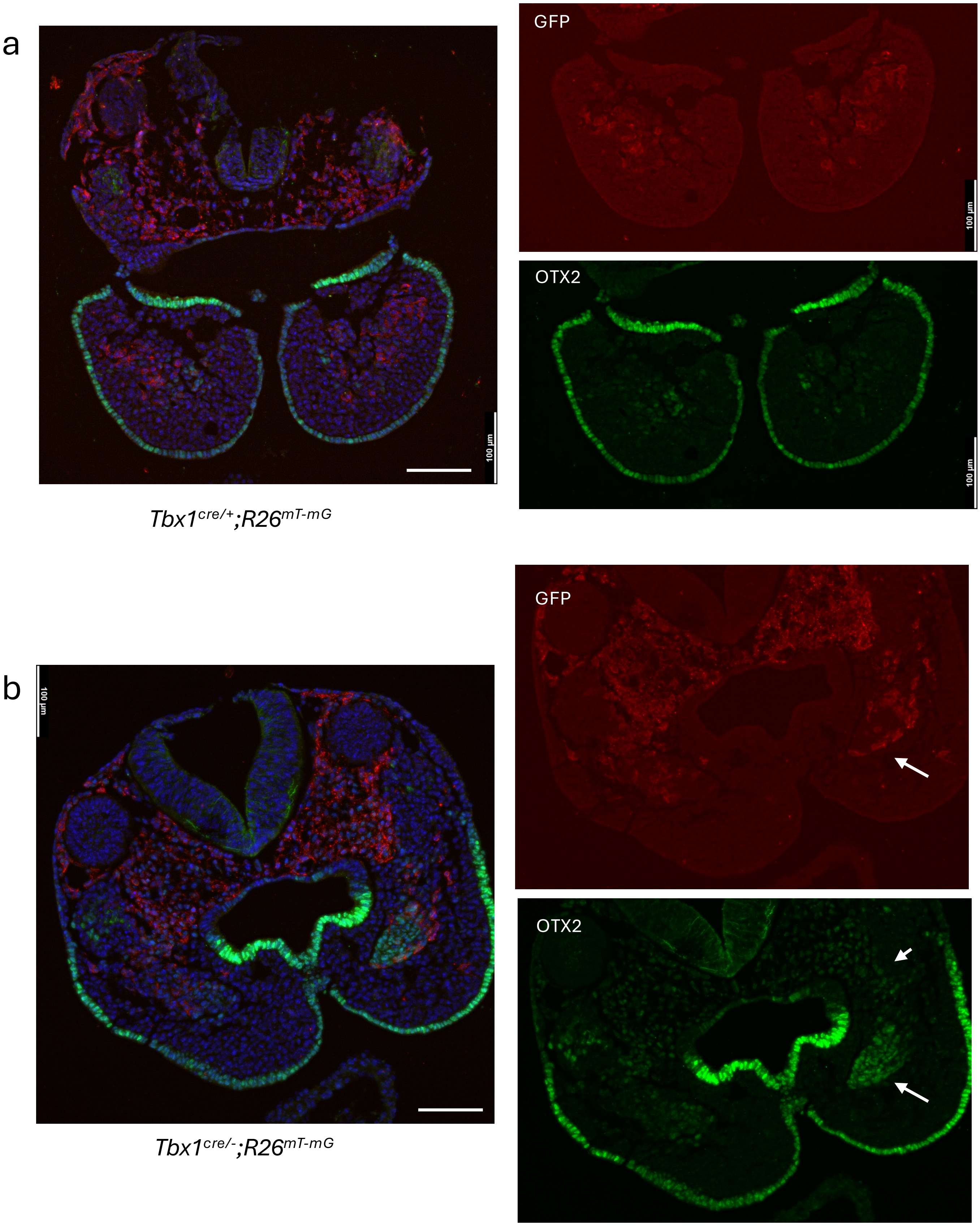


#### Supplementary Figure 12

*Genetic labelling of Tbx1-expressing cells and their descendants reveals partial overlap with ectopic OTX2-expressing cells in homozygous mutant embryos.*

Transverse sections of E9.5 mouse embryos stained with two-colors immunofluorescence anti GFP (genetic label of *Tbx1*-expressing cells, in red) and anti OTX2 (in green): a) control (heterozygous) embryo, merged image (left panel) and split channels green and red (right panels); b) Homozygous mutant embryo, note the region of up regulation in the first pharyngeal arch and head mesenchyme (arrows).

Scale bars are 100 µm.

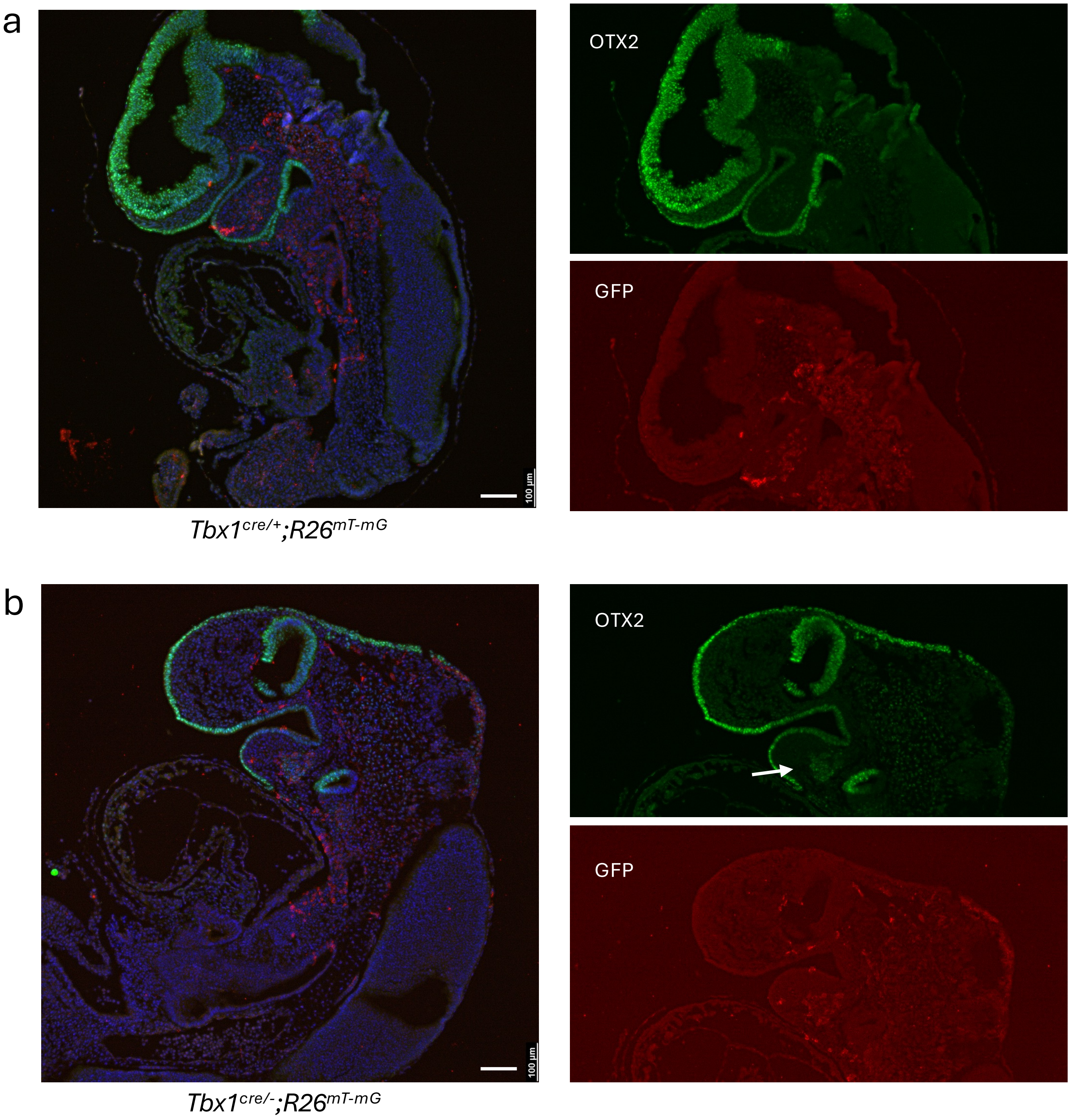


#### Supplementary Figure 13

*Genetic labelling of Tbx1-expressing cells and their descendants reveals partial overlap with ectopic OTX2-expressing cells in homozygous mutant embryos (continued).*

Sagittal sections of E9.5 mouse embryos stained with two-colors immunofluorescence anti GFP (genetic label of *Tbx1*-expressing cells, in red) and anti OTX2 (in green): a) control (heterozygous) embryo, merged image (left panel) and split channels green and red (right panels); b) Homozygous mutant embryo, note the region of up regulation in the first pharyngeal arch (arrow).

Scale bars are 100 µm.
