## Supplementary material for "*Tbx1* stabilizes differentiation of the cardiopharyngeal mesoderm and drives morphogenesis in the pharyngeal apparatus": Suppl. Tab. 3

| **Motif** | **Name** | **q-value** |
| --- | --- | --- |
| 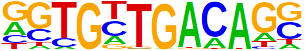 | Tbx20(T-box)/Heart-Tbx20-ChIP-Seq(GSE29636)/Homer | 0.0000 |
| 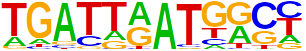 | Hoxb4(Homeobox)/ES-Hoxb4-ChIP-Seq(GSE34014)/Homer | 0.0002 |
| 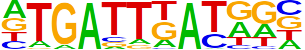 | PBX2(Homeobox)/K562-PBX2-ChIP-Seq(Encode)/Homer | 0.0002 |
| 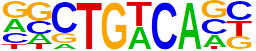 | Meis1(Homeobox)/MastCells-Meis1-ChIP-Seq(GSE48085)/Homer | 0.0003 |
| 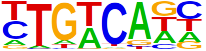 | Tgif1(Homeobox)/mES-Tgif1-ChIP-Seq(GSE55404)/Homer | 0.0015 |
| 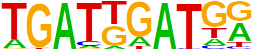 | HOXA1(Homeobox)/mES-Hoxa1-ChIP-Seq(SRP084292)/Homer | 0.0015 |
| 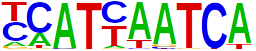 | Pdx1(Homeobox)/Islet-Pdx1-ChIP-Seq(SRA008281)/Homer | 0.0047 |
| 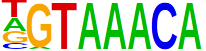 | Foxo3(Forkhead)/U2OS-Foxo3-ChIP-Seq(E-MTAB-2701)/Homer | 0.0060 |
| 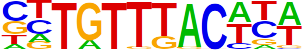 | Foxa2(Forkhead)/Liver-Foxa2-ChIP-Seq(GSE25694)/Homer | 0.0060 |
| 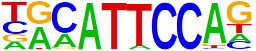 | TEAD3(TEA)/HepG2-TEAD3-ChIP-Seq(Encode)/Homer | 0.0060 |
| 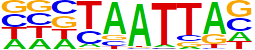 | DLX2(Homeobox)/BasalGanglia-Dlx2-ChIP-seq(GSE124936)/Homer | 0.0060 |
| 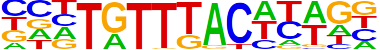 | Foxa3(Forkhead)/Liver-Foxa3-ChIP-Seq(GSE77670)/Homer | 0.0060 |
| 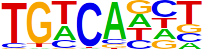 | Tgif2(Homeobox)/mES-Tgif2-ChIP-Seq(GSE55404)/Homer | 0.0197 |
| 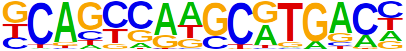 | PAX5(Paired,Homeobox)/GM12878-PAX5-ChIP-Seq(GSE32465)/Homer | 0.0197 |
| 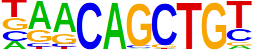 | Myf5(bHLH)/GM-Myf5-ChIP-Seq(GSE24852)/Homer | 0.0197 |
| 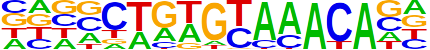 | Fox:Ebox(Forkhead,bHLH)/Panc1-Foxa2-ChIP-Seq(GSE47459)/Homer | 0.01970.0246 |
| 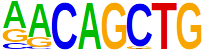 | MyoG(bHLH)/C2C12-MyoG-ChIP-Seq(GSE36024)/Homer | 0.02460.0249 |
| 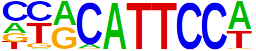 | TEAD1(TEAD)/HepG2-TEAD1-ChIP-Seq(Encode)/Homer | 0.02490.0325 |
| 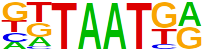 | Nkx6.1(Homeobox)/Islet-Nkx6.1-ChIP-Seq(GSE40975)/Homer | 0.0325 |
| 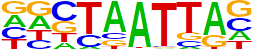 | En1(Homeobox)/SUM149-EN1-ChIP-Seq(GSE120957)/Homer | 0.0325 |
| 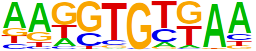 | Tbr1(T-box)/Cortex-Tbr1-ChIP-Seq(GSE71384)/Homer | 0.0325 |
| 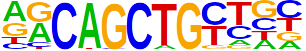 | MyoD(bHLH)/Myotube-MyoD-ChIP-Seq(GSE21614)/Homer | 0.0325 |
| 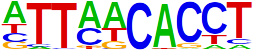 | Eomes(T-box)/H9-Eomes-ChIP-Seq(GSE26097)/Homer | 0.0361 |
| 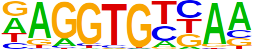 | Tbx6(T-box)/ESC-Tbx6-ChIP-Seq(GSE93524)/Homer | 0.0371 |
| 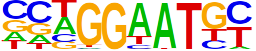 | TEAD4(TEA)/Tropoblast-Tead4-ChIP-Seq(GSE37350)/Homer | 0.0459 |
| 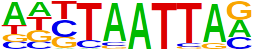 | Lhx3(Homeobox)/Neuron-Lhx3-ChIP-Seq(GSE31456)/Homer | 0.0467 |
| 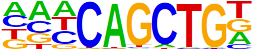 | Ap4(bHLH)/AML-Tfap4-ChIP-Seq(GSE45738)/Homer | 0.0486 |
| 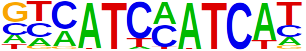 | HOXA2(Homeobox)/mES-Hoxa2-ChIP-Seq(Donaldson_et_al.)/Homer | 0.0494 |
| 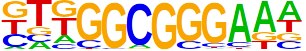 | E2F3(E2F)/MEF-E2F3-ChIP-Seq(GSE71376)/Homer | 0.0494 |
| 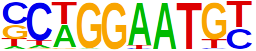 | TEAD2(TEA)/Py2T-Tead2-ChIP-Seq(GSE55709)/Homer | 0.0494 |
